# The ubiquitin ligase HOIL-1L regulates immune responses by interacting with linear ubiquitin chains

**DOI:** 10.1101/2021.04.13.439587

**Authors:** Carlos Gomez-Diaz, Gustav Jonsson, Katrin Schodl, Luiza Deszcz, Annika Bestehorn, Kevin Eislmayr, Jorge Almagro, Anoop Kavirayani, Lilian M Fennell, Astrid Hagelkruys, Pavel Kovarik, Josef M Penninger, Fumiyo Ikeda

## Abstract

The Linear Ubiquitin Assembly Complex (LUBAC), composed of HOIP, HOIL-1L and SHARPIN, promotes Tumor Necrosis Factor (TNF)-dependent NF-κB signaling in diverse cell types. HOIL-1L contains an Npl4 Zinc Finger (NZF) domain that specifically recognizes linear ubiquitin chains, but its physiological role *in vivo* has remained unclear. Here, we demonstrate that the HOIL-1L NZF domain has important regulatory functions in inflammation and immune responses in mice. We generated knockin mice (*Hoil-1l*^*T20;A;R208A/T201A;R208A*^) expressing a HOIL-1L NZF mutant, and observed attenuated responses to TNF- and LPS-induced shock, including prolonged survival, stabilized body temperature, reduced cytokine production and liver damage markers. Cells derived from the HOIL-1L knockin mice show reduced TNF-dependent NF-κB activation and incomplete recruitment of HOIL-1L into TNF Receptor (TNFR) Complex I. We further show that the HOIL-1L-NZF domain cooperates with SHARPIN to prevent TNFR-dependent skin inflammation. Collectively, our data suggest that linear ubiquitin-chain binding by HOIL-1L regulates immune responses and inflammation *in vivo*.

## Introduction

Tumor Necrosis Factor (TNF) is a pleiotropic cytokine involved in the maintenance and regulation of homeostasis and in the response to bacterial and viral infections (Webster and Vucic, 2020; Karki *et al*., 2021). TNF binding to TNF Receptor 1 (TNFR1) leads to the formation of TNFR Complex I and, in turn, Nuclear Factor kappa B (NF-κB)-signaling. This signaling activates cell survival by inducing the transcription of pro-inflammatory and anti-apoptotic genes (Gómez-Díaz and Ikeda, 2019; Peltzer and Walczak, 2019). TNFR Complex I includes, among other proteins, Receptor Interacting serine/threonine Protein Kinase 1 (RIPK1), TNF Receptor type 1 Associated DEATH Domain (TRADD), TNF Receptor Associated Factor 2 (TRAF2) and cellular Inhibitor of Apoptosis Protein (cIAP)1/2 (Peltzer and Walczak, 2019) (Supplementary Figure 1A).

**Figure 1.**
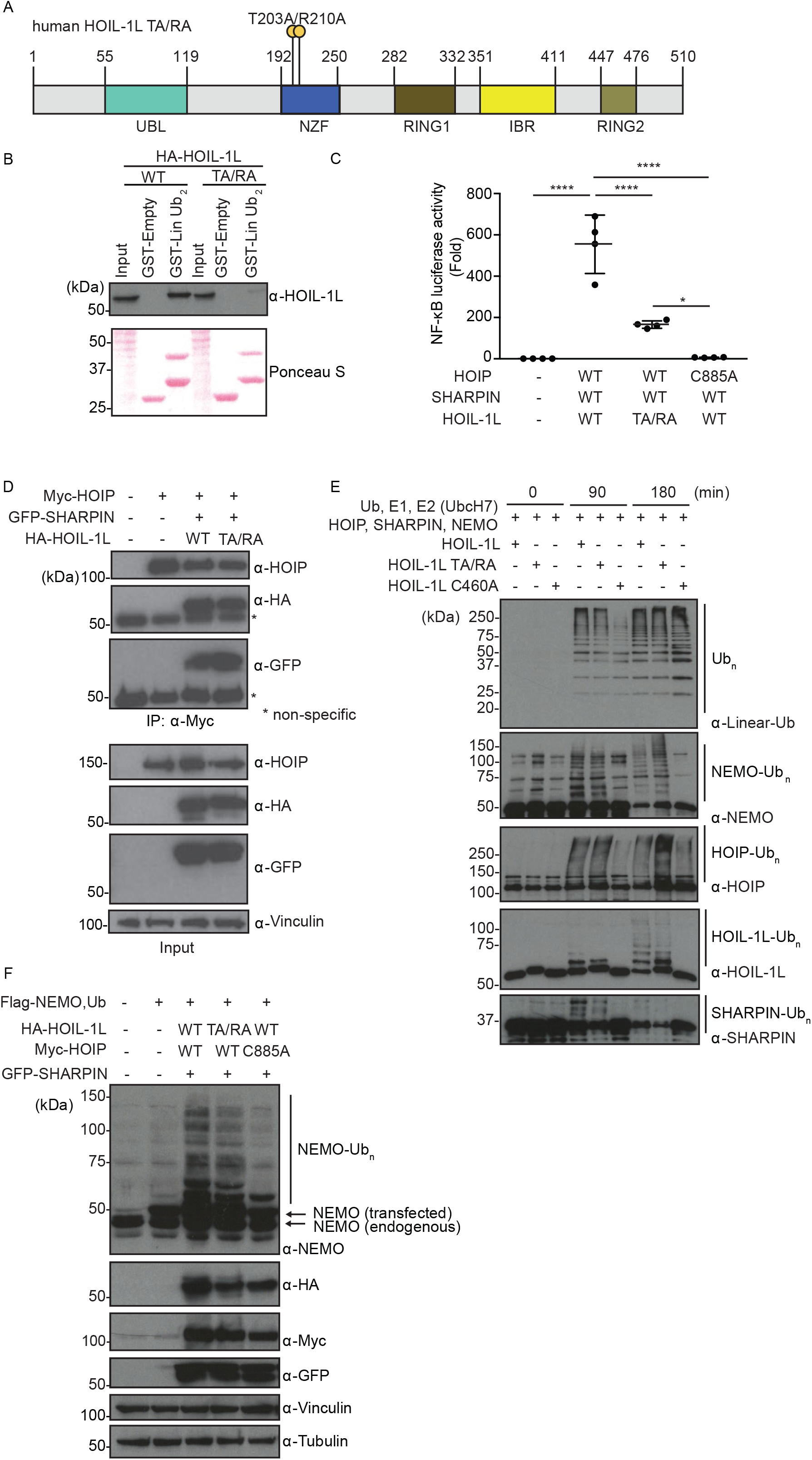
Mutations in the HOIL-1L-NZF (Thr 203 Ala/ Arg 210 Ala) reduce the NF-κB reporter activity without blocking LUBAC catalytic activity. **A**. Schematic representation of human HOIL-1L with the annotated domains and the T203A/R210A double point mutation in the HOIL-1L-NZF domain (human HOIL-1L TA/RA). **B**. GST-pulldown assay examining interaction between linear ubiquitin chain dimer and HOIL-1L (WT or TA/RA). Total cell lysates of HEK293T cells transfected with indicated plasmids were incubated with GST-Empty control or GST-tagged linear di-ubiquitin (GST-Lin Ub_2_). Total cell lysates (Input) and pull-down samples were examined by immunoblotting using α-HOIL-1L. Input amount of total cell lysates and GST-fusion proteins was visualized by Ponceau S staining. **C**. Luciferase-based NF-κB reporter assay using lysates of HEK293T cells co-transfected with an NF-κB reporter plasmid, Renilla-luciferase expression plasmid (internal control), with or without Myc-HOIP (WT or catalytic inactive C885A mutant), Flag-SHARPIN (WT) and HOIL-1L-HA (WT or TA/RA) plasmids. **D**. Immunoblots of co-immunoprecipitation samples examining interaction between Myc-HOIP and HOIL-1L-HA (WT or T203A/R210A) or GFP-SHARPIN. Total cell lysates of HEK293T cells transiently expressing tagged LUBAC components as indicated, were immunoprecipitated by using an α-Myc antibody. Total cell lysates (Input) and immunoprecipitates (IP: α-Myc) were subjected to SDS-PAGE followed by immunoblotting using the indicated antibodies. ***E***. *In vitro* ubiquitination assays using the recombinant proteins of ubiquitin (Ub), mouse Ube1 (E1), E2 (UbcH7), HOIP, HOIL-1L (WT, TA/RA or a mutant of ubiquitin loading site C460A), SHARPIN and NEMO. Samples were subjected to SDS-PAGE. Linear ubiquitin chain formation and modifications of NEMO, HOIP, HOIL-1L and SHARPIN were detected by immunoblotting with the indicated antibodies. **F**. LUBAC-induced ubiquitination of NEMO and linear ubiquitin chain formation in HEK293T cells determined by immunoblotting. Total cell lysates transiently expressing Flag-NEMO, Myc-HOIP (WT or C885A mutant), GFP-SHARPIN, HOIL-1L-HA (WT or T203A/R210A mutant) were subjected to SDS-PAGE followed by immunoblot with the indicated antibodies. α-Vinculin antibody was used to monitor loading amount. **Data information**. Data are representative of at least three independent experiments. (C) Data are represented as mean ±SD, ANOVA, n=4, *p-value ≤0.05, ****p-value ≤0.0001.

Post translational modifications, including ubiquitination and phosphorylation, regulate signal transduction downstream of TNFR complex I.For example, the E3 ubiquitin ligases cIAP1/2 ubiquitinate themselves and RIPK1, leading to the recruitment of the Linear UBiquitin Chain Assembly Complex (LUBAC) (Haas *et al*., 2009) and a kinase complex IKK. LUBAC is the only known mammalian E3 ubiquitin ligase, which generates linear (Met1-linked) ubiquitin chains and stabilizes TNFR Complex I by ubiquitinating various substrates, such as NF-κB Essential Modifier (NEMO) and RIPK1 (Supplementary Figure 1A) (Justus and Ting, 2015; Gómez-Díaz and Ikeda, 2019; Peltzer and Walczak, 2019). Thus, LUBAC promotes NF-κB signaling and cell survival via TNFR Complex I (Gómez-Díaz and Ikeda, 2019).

Sustained or excessive TNF stimulation leads to dissociation of RIPK1 from TNFR Complex I and its association with TNFR Complex II (Supplementary Figure 1A) (Justus and Ting, 2015; Witt and Vucic, 2017). TNFR Complex II induces either apoptosis, by activating caspase-8, followed by caspase-3 (Witt and Vucic, 2017), or necroptosis by activating RIPK3 and Mixed Lineage Kinase domain Like pseudo kinase (MLKL) (Witt and Vucic, 2017). LUBAC-mediated linear ubiquitination negatively regulates TNFR Complex II -dependent apoptosis (Asaoka and Ikeda, 2015; Sasaki and Iwai, 2015; Peltzer and Walczak, 2019).

LUBAC consists of three proteins: two RING-in between-RING (RBR)-E3 ubiquitin ligases, the Heme-Oxidized IRP2 ubiquitin Ligase 1 (HOIL-1L) and the HOIL-1L Interacting Protein (HOIP) and the adaptor the Shank-Associated RH Domain-Interacting Protein (SHARPIN) (Gerlach *et al*., 2011; Ikeda *et al*., 2011; Tokunaga *et al*., 2011). Genetic deletion of HOIP or HOIL-1L in mice leads to embryonic lethality due to aberrant endothelial cell death, namely apoptosis and necroptosis in the vasculature (Peltzer *et al*., 2014, 2018). SHARPIN-deficient mice (*Sharpin*^*cpdm/cpdm*^) are viable and display an inflammatory phenotype characterized by chronic proliferative dermatitis (Seymour *et al*., 2007), which is largely resolved by loss of TNF or TNFR (Gerlach *et al*., 2011; Kumari *et al*., 2014; Rickard *et al*., 2014). Mutations in *HOIL-1L* and *HOIP* have been associated with autoimmune disorders, implicating LUBAC in the regulation of immune responses in humans (Boisson *et al*., 2012, 2015).

HOIL-1L is required for proper inflammatory responses and embryonic development (Peltzer *et al*., 2018; Kelsall *et al*., 2019; Fuseya *et al*., 2020). It consists of multiple domains including the Npl4 Zinc Finger (NZF), the ubiquitin-like domain (UBL) and the RBR domain (Supplementary Figure 1B). The biochemical properties of each domain are known: NZF binds linear ubiquitin chains (Sato *et al*., 2011; Fennell, Rahighi and Ikeda, 2018), UBL binds other LUBAC components, and RBR catalyzes ester-bond linkage of ubiquitination (Kelsall *et al*., 2019; Carvajal *et al*., 2020). Threonine 203 and Arginine 210 within the human HOIL-NZF domain are essential to establish the interaction with a linear di-ubiquitin chain (Sato *et al*., 2011). Mutation of these residues results in decreased LUBAC-induced NF-κB reporter activation (Sato *et al*., 2011), and *Hoil-1l*^*-/-*^ Mouse Embryonic Fibroblasts (MEFs) reconstituted with a HOIL-1L-NZF mutant (T201A/R208A) show inefficient recruitment of HOIP and SHARPIN to TNFR Complex I and reduced linear ubiquitination in the TNFR Complex I (Peltzer *et al*., 2018). However, it remains unclear whether the HOIL-1L NZF domain directly regulates the ubiquitin ligase activity of LUBAC and to what extent it controls immunity and inflammation *in vivo*.

## Results

### HOIL-1L NZF mutations block linear ubiquitin chain binding and NF-κB activation without disrupting LUBAC activity

The NZF domain of HOIL-1L (aa 192-250) specifically recognizes linear ubiquitin chains via two key residues, Threonine 203 and Arginine 210 (Figure 1A) (Sato *et al*., 2011). To confirm the importance of these residues for binding, we expressed HOIL-1L wild type (WT) or HOIL-1L T203A/R210A in HEK293T cells and performed pull down assays with GST-linear-di-ubiquitin (GST-Lin Ub_2_). As expected, WT HOIL-1L but not HOIL-1L T203A/R210A interacted with GST-Lin Ub_2_ (Figure 1B).

To verify that the HOIL-1L NZF domain promotes NF-κB activation via LUBAC, we employed a luciferase-based NF-κB reporter assay in HEK293T cells co-expressing catalytically active HOIP. HOIL-1L WT activated the NF-κB reporter with or without SHARPIN, as previously shown (Tokunaga *et al*., 2009; Gerlach *et al*., 2011; Ikeda *et al*., 2011) (Figure 1B, 1C and Supplementary Figure 1B-G). In contrast, NF-κB reporter activity was significantly decreased in cells expressing either HOIL-1L T203A/R210A or HOIL-1L-ΔNZF (Figure 1C, Supplementary Figure 1B-D), even though HOIL-1L T203A/R210A supported LUBAC formation (Figure 1D). We also examined a HOIL-1L-RBR-NZF mutant and the catalytic inactive mutant HOIL-1L C460A (Supplementary Figure 1B). NF-κB reporter activity was significantly decreased with HOIL-1L-RBR-NZF, but increased with HOIL-1L C460A (Supplementary Figure 1D), consistent with previous observations ((Elton *et al*., 2016).

To assess the catalytic activity of LUBAC containing HOIL-1L T203A/R210A, we examined the formation of unanchored linear ubiquitin chains as well as the ubiquitination of known substrates by recombinant proteins *in vitro* (Figure 1E and Supplementary Figure 1H). Similar to HOIL-1L WT, LUBAC containing HOIL-1L T203A/R210A supported the formation of unanchored linear ubiquitin chains (Figure 1E), and the ubiquitination of NEMO, HOIP, HOIL-1L and SHARPIN (Figure 1E). In addition, LUBAC containing HOIL-1L T203A/R210A supported NEMO ubiquitination in HEK293T cells (Figure 1F). Collectively, these results suggest that loss of the HOIL-1L NZF domain impairs LUBAC-dependent activation of the NF-κB pathway without severely compromising LUBAC ligase activity.

Notably, in the absence of SHARPIN, unanchored and anchored ubiquitin chain formation were clearly delayed with HOIL-1L T203A/R210A compared to HOIL-1L WT (Supplementary Figure 1H), suggesting that SHARPIN and the HOIL-1L NZF domain have a collaborative role in promoting the E3 ligase functions of LUBAC.

In summary, our data suggest that the reduction of NF-κB reporter activity by the HOIL-1L NZF mutations are not due to loss of LUBAC activity.

### The HOIL-1L NZF domain is required for endogenous TNF-induced NF-κB signaling

To determine if the HOIL-1L NZF domain and binding linear ubiquitin regulates immune signaling cascades *in vivo*, we generated *Hoil-1l*^*T201A;R208A/T201A;R208A*^ knockin mice (mutations at the equivalent residues to T203 and R210 in human HOIL-1L) using CRSIPR-Cas9 technology (Supplementary Figure 2A-C). *Hoil-1l*^*T201A;R208A/T201A;R208A*^ (referred as *Hoil-1l*^*nzf*/nzf\**^) mice were born at nearly the expected ratio and showed no obvious developmental defects (Figure 2A-B). Histological analysis of various tissues (spleen, liver, intestine and skin) from 11-week-old male mice displayed no clear differences between *Hoil-1l*^*+/+*^ (wild type) and *Hoil-1l*^*nzf*/nzf\**^ further supporting that these mice develop normally under basal conditions (Figure 2C).

**Figure 2.**
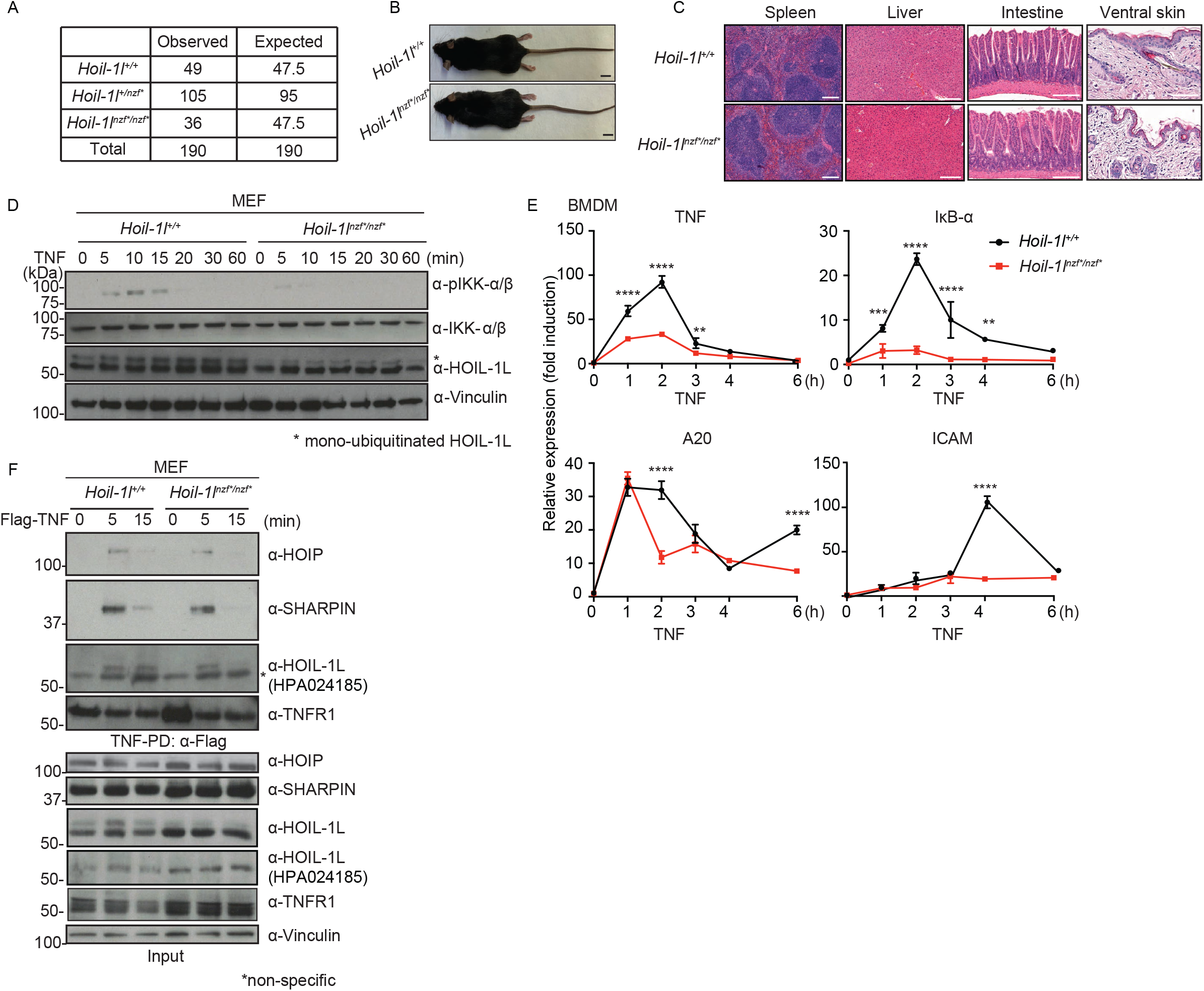
*Hoil-1l*^*nzf*/nzf\**^ knockin mice display no developmental phenotype yet TNF-induced NF-κB activation is reduced in *HOIL-1L*^*NZF*/NZF\**^ cells. **A**. Numbers of weaned and expected mice of the indicated genotypes from *Hoil-1l*^*+/nzf\**^crosses. **B**. Gross appearance image of 11-week-old *Hoil-1l*^+/+^ and *Hoil-1l*^*nzf*/nzf\**^ mice. Scale bar: 10mm. **C**. H&E staining of the indicated tissues from 11-week-old *Hoil-1l*^*+/+*^ and *Hoil-1l*^*nzf*/nzf\**^ mice. Scale bar: 200μm. **D**. Immunoblotting to detect phosphorylation of IKK-α/β in *Hoil-1l*^*+/+*^ and *Hoil-1l*^*nzf*/nzf\**^ MEFs treated with mouse TNF (20 ng/ml) for the indicated time. α-Vinculin antibody was used to monitor loading amount. * indicates mono-ubiquitinated HOIL-1L. **E**. mRNA transcript levels of NF-κB targets (mouse TNF, IκB-α, A20 and ICAM) determined by qRT-PCR in *Hoil-1l*^*+/+*^ and *Hoil-1l*^*nzf*/nzf\**^ BMDMs treated with mouse TNF (20 ng/ml) for the indicated time. Normalization was done to β-actin. **F**. Immunoblotting to detect TNFR Complex I formation in *Hoil-1l*^*+/+*^ and *Hoil-1l*^*nzf*/nzf\**^ MEFs. Total cell lysates (Input) of cells treated with or without human TNF (100 ng/ml), and immunoprecipitates (IP: α-FLAG) were subjected to SDS-PAGE. Recruitment of HOIP, SHARPIN and HOIL-1L was monitored by immunoblotting. α-HOIL-1L (Atlas Antibody, HPA 024185) was used for detecting HOIL-1L in TNFR Complex I and input. α-HOIL-1L (Merck Millipore, MABC576) was additionally used to detect HOIL-1L in input. α-Vinculin antibody was used to monitor loading amount. **Data information**. Data are representative of at least three independent experiments. (E) Data are represented as mean ±SD. ANOVA, n=3, **p-value ≤0.01, ***p-value ≤0.001, ****p-value ≤0.0001.

To assess the physiological role of the HOIL-1L NZF domain in NF-κB signaling, we established immortalized MEFs from *Hoil-1l*^*nzf*/nzf\**^ mice and exposed them to TNF. *Hoil-1l*^*nzf*/nzf\**^ MEFs displayed similar levels of unmodified HOIL-1L but lower levels of mono-ubiquitinated HOIL-1L (Kelsall *et al*., 2019) compared to wild-type MEFs (Figure 2D), suggesting that the HOIL-1L NZF domain and binding linear ubiquitin chains contribute to HOIL-1L mono-ubiquitination in cells. Importantly, TNF-induced phosphorylation of the NF-κB signaling component IKK-α/β was reduced in *Hoil-1l*^*nzf*/nzf\**^ MEFs compared to wild type MEFs (Figure 2D). Moreover, the TNF-induced expression of four NF-κB-target genes was reduced in both *Hoil-1l*^*nzf*/nzf\**^ MEFs (Supplementary Figure 2D) and *Hoil-1l*^*nzf*/nzf\**^ bone marrow-derived macrophages (BMDMs) (Figure 2E). These data indicate that loss of the HOIL-1L NZF domain function inhibits TNF-induced NF-κB signaling in non-immune and immune cells.

To decipher how the HOIL-1L NZF domain regulates the TNF-induced NF-κB pathway, we stimulated MEFs with Flag-TNF and performed pull-down assays with anti-Flag to analyze TNFR Complex I formation. Wild type and *Hoil-1l*^*nzf*/nzf\**^ MEFs expressed similar levels of the individual LUBAC components (Figure 2F, input) and showed comparable TNF-induced recruitment and modification of RIPK1 in the TNFR complex I (Supplementary Figure 2E). In contrast, recruitment of HOIP, SHARPIN and HOIL-1L into the TNFR complex I were reduced and more transient in *Hoil-1l*^*nzf*/nzf\**^ MEFs (Figure 2F). These data suggest that diminished recruitment of LUBAC to TNFR Complex I could underlie the reduced NF-κB activation in *Hoil-1l*^*nzf*/nzf\**^ MEFs.

Next, we examined whether the HOIL-1L NZF domain participates in LUBAC’s negative regulation of TNF-induced apoptosis (Asaoka and Ikeda, 2015; Sasaki and Iwai, 2015; Peltzer and Walczak, 2019). Wild type and *Hoil-1l*^*nzf*/nzf\**^ MEFs treated with TNF and cycloheximide (CHX) showed similar caspase-8 activity (Supplementary Figure 2F), and similar levels of cleaved caspase-3 (CASP3) and cleaved poly ADP ribose polymerase (PARP) (Supplementary Figure 2G), suggesting that the HOIL-1L NZF domain is not required for LUBAC to inhibit apoptosis induction in these cells.

Collectively, these results suggest that the HOIL-1L NZF domain is dispensable for LUBAC’s inhibition of TNF-induced apoptosis, but promotes TNF-induced NF-κB signaling, likely by maintaining LUBAC components in the TNFR Complex I.

### HOIL-1L NZF mutations attenuate TNF-induced shock and LPS-induced septic shock in mice

Having determined that the HOIL-1L NZF domain regulates the TNF-induced NF-κB pathway in cells, we sought to address its contribution *in vivo* during TNF-induced shock. To this end, mouse TNF (mTNF) was intravenously injected into wild type and *Hoil-1l*^*nzf*/nzf\**^ mice and their responses were monitored for up to 12 hours. Wild type mice displayed a drop in body surface temperature starting at 6 hours post-TNF injection, as expected, whereas *Hoil-1l*^*nzf*/nzf\**^ mice were more resistant (Figure 3A). We investigated the TNF-induced immune responses in these mice by measuring cytokine levels in serum. The TNF-induced pro-inflammatory cytokines Interleukin (IL-6), IL-12 (p70) and Granulocyte Colony Stimulating Factor (G-CSF), which are all NF-κB targets (Pahl, 1999), were lower in serum from *Hoil-1l*^*nzf*/nzf\**^ mice than wild type mice (Figure 3B-D). Overall, these data suggest that *Hoil-1l*^*nzf*/nzf\**^ mice have lower TNF-induced NF-κB signaling and are thus more resistant to TNF-induced shock.

**Figure 3.**
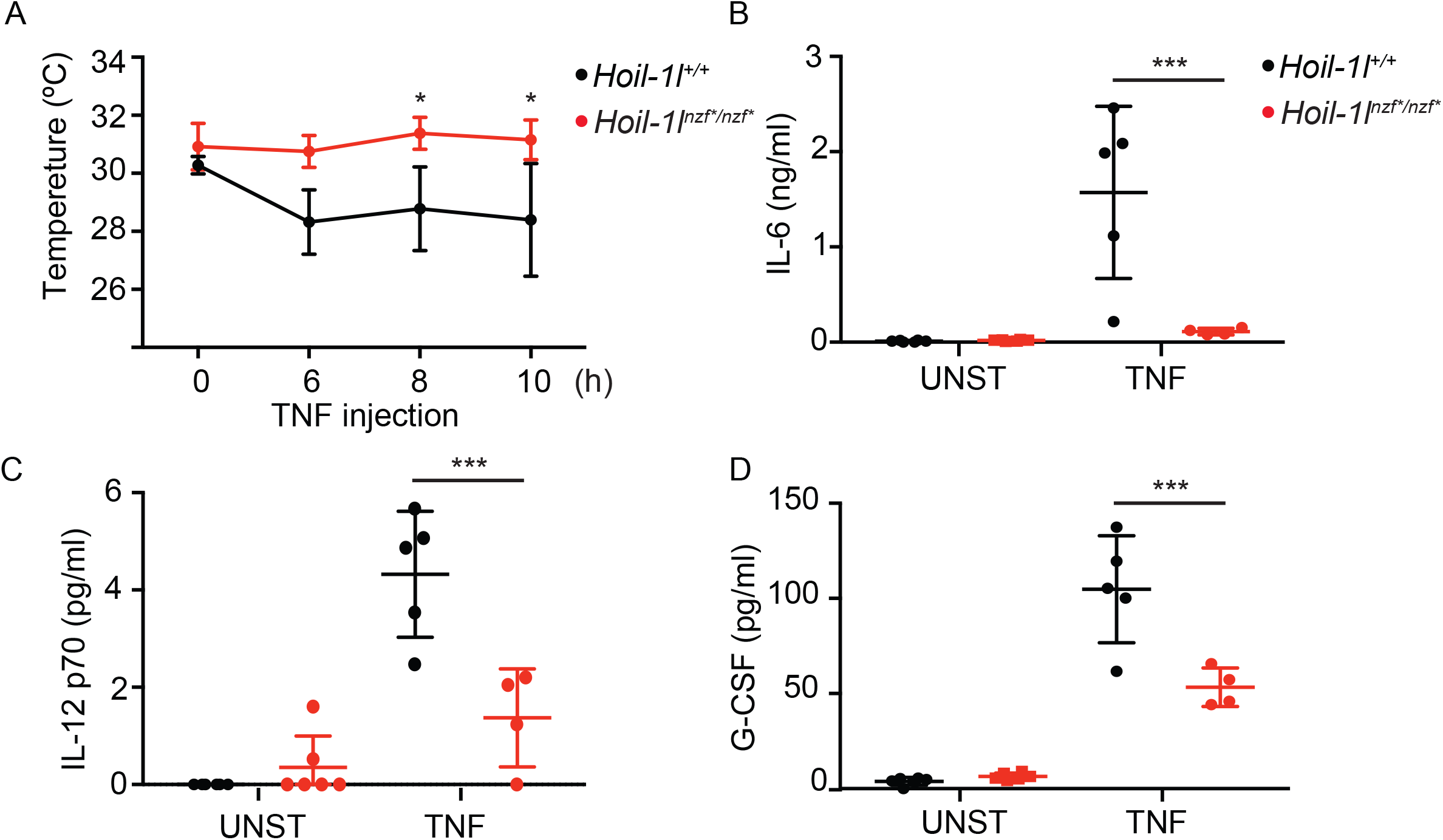
*Hoil-1l*^*nzf*/nzf\**^ mice show reduced responses to TNF-induced shock. **A**. Temperature of *Hoil-1l*^*+/+*^ (n=5) and *Hoil-1l*^*nzf*/nzf\**^ (n=4) upon intravenous TNF injection (450μg/kg) monitored over time. (n=5), *Hoil-1l* **B-D**. IL-6 (B), IL-12 (C) and G-CSF (D) levels in the serum of unstimulated (UNST) and TNF-injected mice. After 12 hours of TNF injection, serum levels of indicted cytokines were measured by ELISA. UNST: *Hoil-1l*^*+/+*^ (n=3), *Hoil-1l*^*nzf*/nzf\**^ (n=3), TNF: *Hoil-1l*^*+/+*^ (n=5), *Hoil-1l*^*NZF*/NZF\**^ (n=4). **Data information**. Data are represented as mean ±SD. ANOVA, *p-value ≤0.05, ***p-value ≤0.001.

To further address the role of the HOIL-1L NZF domain in immune responses, we examined LPS-induced septic shock, which is largely dependent on the TNF signaling pathway (Mandal *et al*., 2018; Vandewalle *et al*., 2019). Intriguingly, *Hoil-1l*^*nzf*/nzf\**^ mice survived longer than wild type littermates after LPS-induced septic shock (Figure 4A), despite similarly sharp drops in temperature (Figure 4B), similar levels of infiltrating myeloid cells in the peritoneal cavity (Supplementary Figure 3A-D), and similar serum levels of the pro-inflammatory cytokines, IL-6, IL-27 and G-CSF (Figure 4D-F). LPS-dependent induction of four NF-κB target genes was also comparable in wild type and *Hoil-1l*^*nzf*/nzf\**^ BMDM, suggesting that HOIL-1L NZF is not directly involved in the LPS-TLR4-NF-κB pathway (Figure 4C). However, the serum levels of TNF, IL-1α and IL-1β were decreased in *Hoil-1l*^*nzf*/nzf\**^mice compared to wild type mice (Figure 4G-I) suggesting that these cytokines could underlie the distinct LPS-induced immune responses. Furthermore, *Hoil-1l*^*nzf*/nzf\**^ mice displayed decreased serum levels of the liver enzyme aspartate aminotransferase (AST) compared to wild type after LPS injection, indicating reduced liver damage markers in *Hoil-1l*^*nzf*/nzf\**^ mice (Figure 4J-K). These results indicate that the HOIL-1L NZF domain promotes LPS-induced septic shock, accompanied with selected cytokine induction and increased levels of liver damage markers.

**Figure 4.**
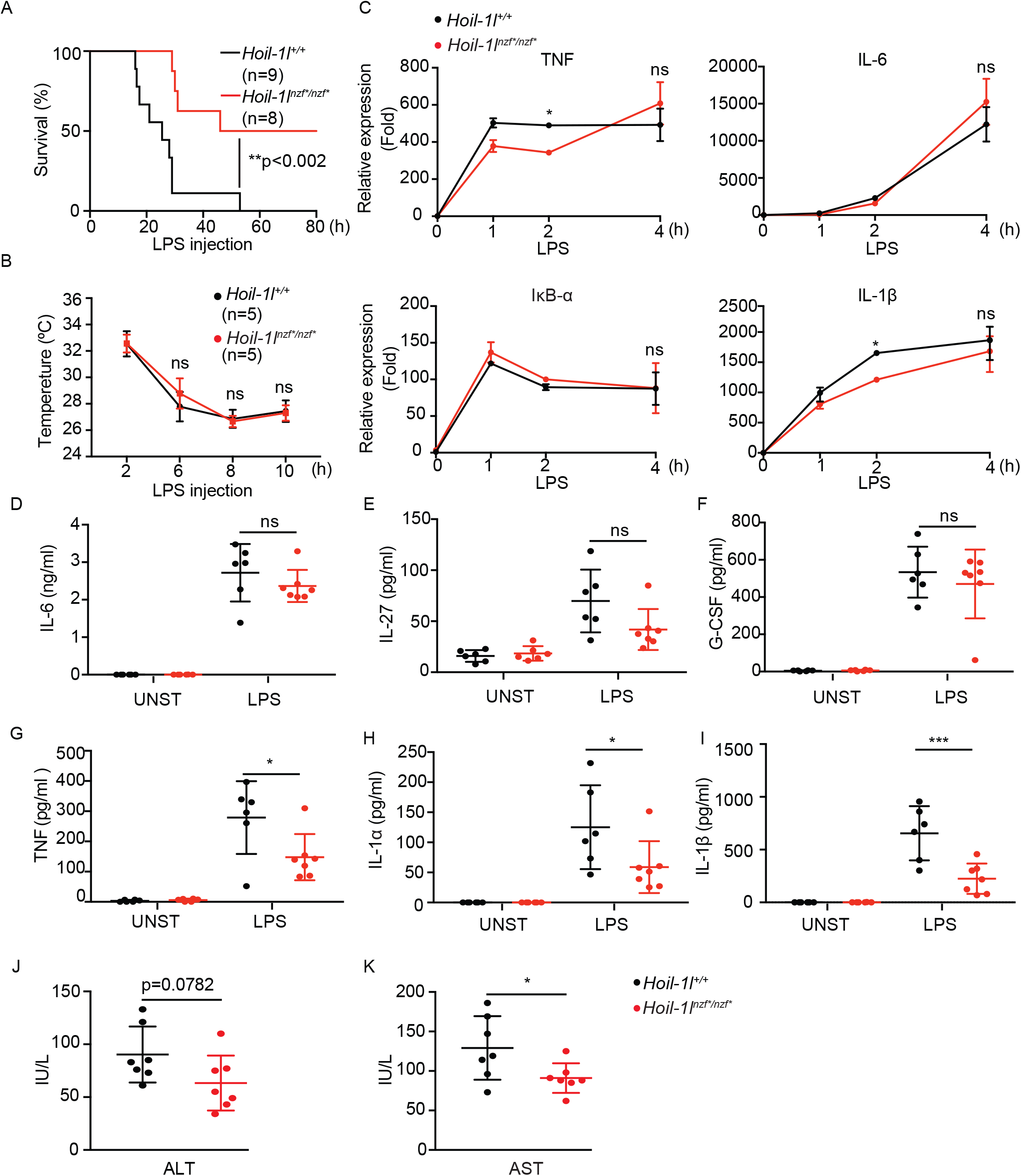
*Hoil-1l*^*nzf*/nzf\**^mice are more resistant to LPS-induced septic shock than wild type mice. **A**.Survival curve of *Hoil-1l*^*+/+*^ (n=9) and *Hoil-1l*^*nzf*/nzf\**^ (n=8) mice upon LPS injection. *Hoil-1l*^*+/+*^ (n=9) and *Hoil-1l* ^*NZF*/NZF\**^ (n=8) mice were injected with 30 mg/kg of LPS and the survival was monitored up to 80 hours. **B**.Body temperature of *Hoil-1l*^*+/+*^ (n=5) and *Hoil-1l*^*nzf*/nzf\**^ (n=5) mice upon LPS injection (30mg/kg) monitored over time. **C**.qRT-PCR to monitor mRNA transcript levels of TNF, IL-6, IκB-α, IL-1β determined in *Hoil-1l*^*+/+*^ and *Hoil-1l*^*nzf*/nzf\**^ BMDMs treated with LPS (10 ng/ml) for the indicated time. Normalization was done to β-actin. **D-I**. ELISA to monitor IL-27, GCSF, IL-6, TNF, IL-1α and IL-1β levels in the serum of control and LPS-injected (30mg/kg) mice. Samples for LPS-injected mice were collected after 12 hours of LPS injection. Unstimulated (UNST): *Hoil-1l*^*+/+*^ (n=3) and *Hoil-1l*^*nzf*/nzf\**^ (n=3), and LPS-injected (LPS): *Hoil-1l*^*+/+*^ (n=6) and *Hoil-1l*^*nzf*/nzf\**^ (n=7). **J-K**. ALT and AST levels in the sera of *Hoil-1l*^*+/+*^ (n=7) and *Hoil-1l*^*nzf*/nzf\**^ (n=7) mice 12 hours post LPS injection (30mg/kg). **Data information**. (A) Log-rank (Mantel-Cox test) **p-value=0.0018. (B-I) Data are represented as mean ±SD, ANOVA. (C) Representative data from three independent experiments, n=3, *p-value ≤0.05. (D-I) *p-value ≤0.05, ***p-value ≤0.001. (J-K) Data are presented as mean ±SD, Students-t, *p-value ≤0.05.

We infer that mutations in the HOIL-1L NZF domain reduce TNF-induced NF-κB signaling and render mice resistant to TNF- and LPS-induced shock.

### HOIL-1L NZF mutations exacerbate systemic inflammation in *Sharpin*^*cpdm/cpdm*^ mice

Given the redundant biochemical properties of HOIL-1L and SHARPIN *in vitro* (Supplementary Figure 1H), we investigated redundancy *in vivo* by generating *Hoil-1l*^*nzf*/nzf\**^ mice in a SHARPIN-deficient background (*Sharpin*^*cpdm/cpdm*^). *Hoil-1l*^*nzf*/nzf\**^; *Sharpin*^*cpdm/cpdm*^ mice were viable and born at nearly the Mendelian ratio (Figure 5A), but presented visible skin inflammation already at 2 weeks after birth, appeared smaller, and had reduced body weight compared to all other littermates, including *Sharpin*^*cpdm/cpdm*^ mice (Fig 5B-C). At 4 weeks after birth, *Hoil-1l*^*nzf*/nzf\**^; *Sharpin*^*cpdm/cpdm*^ mice displayed systemic inflammation characterized by inflammatory cell infiltration in the liver, intestines and lung (Supplementary Figure 4A).

**Figure 5.**
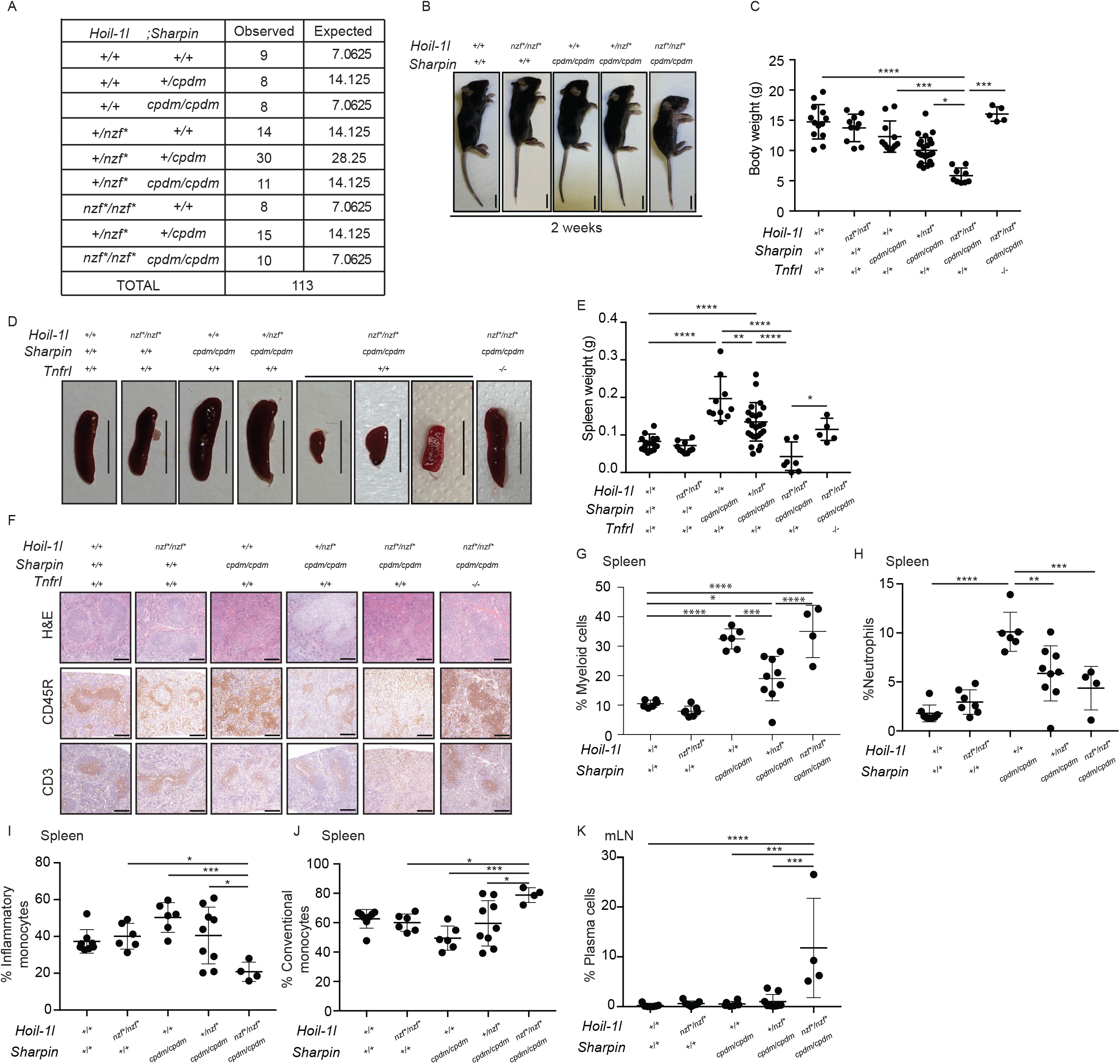
*Hoil-1l*^*nzf*/nzf\**^; *Sharpin*^*cpdm/cpdm*^ mice show systemic inflammation and a distinctive immune cell composition. **A**.Numbers of weaned and expected mice of the indicated genotypes from *Hoil-1l*^*+/nzf\**^; *Sharpin*^*+/cpdm*^ crosses. **B**.Representative image of the gross appearance from 2-week-old mice of the indicated genotypes. Scale bar: 10 mm. **C**.Body weight of 4-week-old mice from the indicated genotypes. **D**.Representative images of the spleen of indicated genotype. **E**.Spleen weight of 4-week-old mice from the indicated genotypes. **F**.H&E staining, CD45R and CD3 immunostaining of spleen sections from 4-week-old mice of the indicated genotypes. Scale bar: 50 μm. **G**.Percentage of viable myeloid cells (CD11b^+^), normalized to the frequency of viable CD45^+^ cells, in the spleen from 4-week-old mice of the indicated genotypes. **H**.Percentage of neutrophils (CD11b^Hi^ Ly6G^+^) out of viable CD45^+^ cells in the spleen from 4-week-old mice of the indicated genotypes. **I-J**. Percentage of inflammatory monocytes (CD11c^-^ CD11b^+^ Ly6G^-^ Ly6C^+^) (G) and conventional monocytes (CD11c^-^ CD11b^+^ Ly6G^-^ Ly6C^-^) (H), out of viable CD45^+^ CD11b^+^ cells, in the spleen from 4-week-old mice of the indicated genotypes. **K**. Percentage of plasma cells (CD138^+^ CD28^+^), out of viable B220^+^ Lin^-^ cells, in the mesenteric lymph node (mLN) from 4-week-old mice of the indicated genotypes. The lineage (Lin) cocktail consists of TCRβ, DX5, CD11b and CD23. **Data information**. Data are represented as mean ±SD. ANOVA, **p-value≤0.01, ***p-value≤0.001, ****p-value ≤0.0001. (G-K, L) *Hoil-1l*^*+/+*^; *Sharpin*^*+/+*^ (n=6), *Hoil-1l*^*nzf*/nzf\**^; *Sharpin*^*+/+*^ (n=7), *Hoil-1l*^*+/+*^; *Sharpin*^*cpdm/cpdm*^ (n=6), *Hoil-1l*^*+/nzf\**^; *Sharpin*^*cpdm/cpdm*^ (n=9), *Hoil-1l*^*nzf*/nzf\**^; *Sharpin*^*cpdm/cpdm*^ (n=4).

*Hoil-1l*^*nzf*/nzf\**^; *Sharpin*^*cpdm/cpdm*^ mice had smaller spleens compared to all other littermates (Figure 5D-E). However, histopathological analysis revealed that disruption of the HOIL-1L NZF did not exacerbate the disruption of the splenic structure in SHARPIN-deficient mice (Figure 5F, H&E). Mutation of the HOIL-1L NZF domain did not exacerbate the reduced frequency of CD45R^+^CD19^+^ B cells nor the increased frequency of myeloid cells in *Sharpin*^*cpdm/cpdm*^ spleens (Supplementary Figure 4B, Figure 5G, Supplementary Figure 4E) (Gurung, Sharma and Kanneganti, 2016; Sharma *et al*., 2019). Although CD45R/B220^+^ B cells and CD3^+^ T cells were present in *Hoil-1l*^*nzf*/nzf\**^; *Sharpin*^*cpdm/cpdm*^ spleens, their distributions were altered compared to spleens of littermates (Figure 5F). The frequency of mature B cells and plasma cells in the spleen were also similar in all analyzed genotypes (Supplementary Figure 4C-D). Neutrophils and inflammatory monocytes were reduced in the spleens of *Hoil-1l*^*nzf*/nzf\**^; *Sharpin*^*cpdm/cpdm*^ mice compared to *Sharpin*^*cpdm/cpdm*^ mice (Figure 5H-I, Supplementary Figure 4F-G) whereas conventional monocytes were increased (Figure 5J, Supplementary Figure 4F). The percentage of total T cells in the spleen was similar across the genotypes analyzed (Supplementary Figure 4H), as was the percentage of CD4^+^ and CD8^+^ T cells (Supplementary Figure 4I-J). Collectively, our results show that mutation of the HOIL-1L NZF domain alters the composition of the myeloid cell compartment in the spleens of SHARPIN-deficient mice.

SHARPIN deficiency, irrespective of the HOIL-1L NZF domain, increased the frequency of myeloid cells (Supplementary Figure 5A) and decreased the frequency of B cells (Supplementary Figure 5B) in the mesenteric lymph node (mLN), without affecting the frequency of mature B cells (Supplementary Figure 5C). Strikingly, only *Hoil-1l*^*nzf*/nzf\**^; *Sharpin*^*cpdm/cpdm*^ mice displayed an increase in plasma cells in the mesenteric lymph nodes, revealing clear additive effects of HOIL-1L NZF and SHARPIN in plasma cell responses (Figure 5K, Supplementary Figure 5D).

Collectively, these results reveal profound differences in the immune cell composition of *Hoil-1l*^*nzf*/nzf\**^; *Sharpin*^*cpdm/cpdm*^ mice compared to SHARPIN-deficient mice and suggest a different inflammatory process in these mice that could underlie their more severe phenotype.

### TNFR1 knockout resolves the inflammatory skin phenotype of *Hoil-1l*^*nzf*/nzf\**^; *Sharpin* ^*cpdm/cpdm*^ mice

The main inflammatory phenotype in SHARPIN-deficient mice is chronic proliferative dermatitis accompanied by keratinocyte apoptosis (Gerlach *et al*., 2011; Ikeda *et al*., 2011; Kumari *et al*., 2014; Rickard *et al*., 2014). We analyzed skin sections derived from *Hoil-1l*^*nzf*/nzf\**^; *Sharpin*^*cpdm/cpdm*^ mice and observed thickening of the keratin layer (H&E and Keratin 14 (KRT14)), apoptotic cells (cleaved caspase-3) and neutrophil infiltration (Lys6G) at 2-weeks of age (Figure 6A) and at 4-weeks of age (Figure 6B-C). These inflammatory characteristics were not observed in any of the other genotypes (Figure 6A-C). At 4-weeks of age, dermatitis was observed even with a single HOIL-1L NZF mutant allele in the *Sharpin*^*cpdm/cpdm*^ background (Figure 6B-C). Loss of HOIP in mice converts the cell death-driven dermatitis in SHARPIN-deficient mice to T cell-predominant autoimmune lesions (Sasaki *et al*., 2019). Therefore, we investigated whether T-cell infiltration in the skin of *Sharpin*^*cpdm/cpdm*^ mice was increased by the HOIL-1L NZF mutations. Indeed, the number of CD3-positive T cells in the skin was increased in *Hoil-1l*^*nzf*/nzf\**^; *Sharpin*^*cpdm/cpdm*^ mice when compared to *Sharpin*^*cpdm/cpdm*^ mice (Figure 6D-E), suggesting that HOIL-1L NZF cooperates with SHARPIN to restrict inflammation displaying characteristics of autoimmunity-like processes.

**Figure 6.**
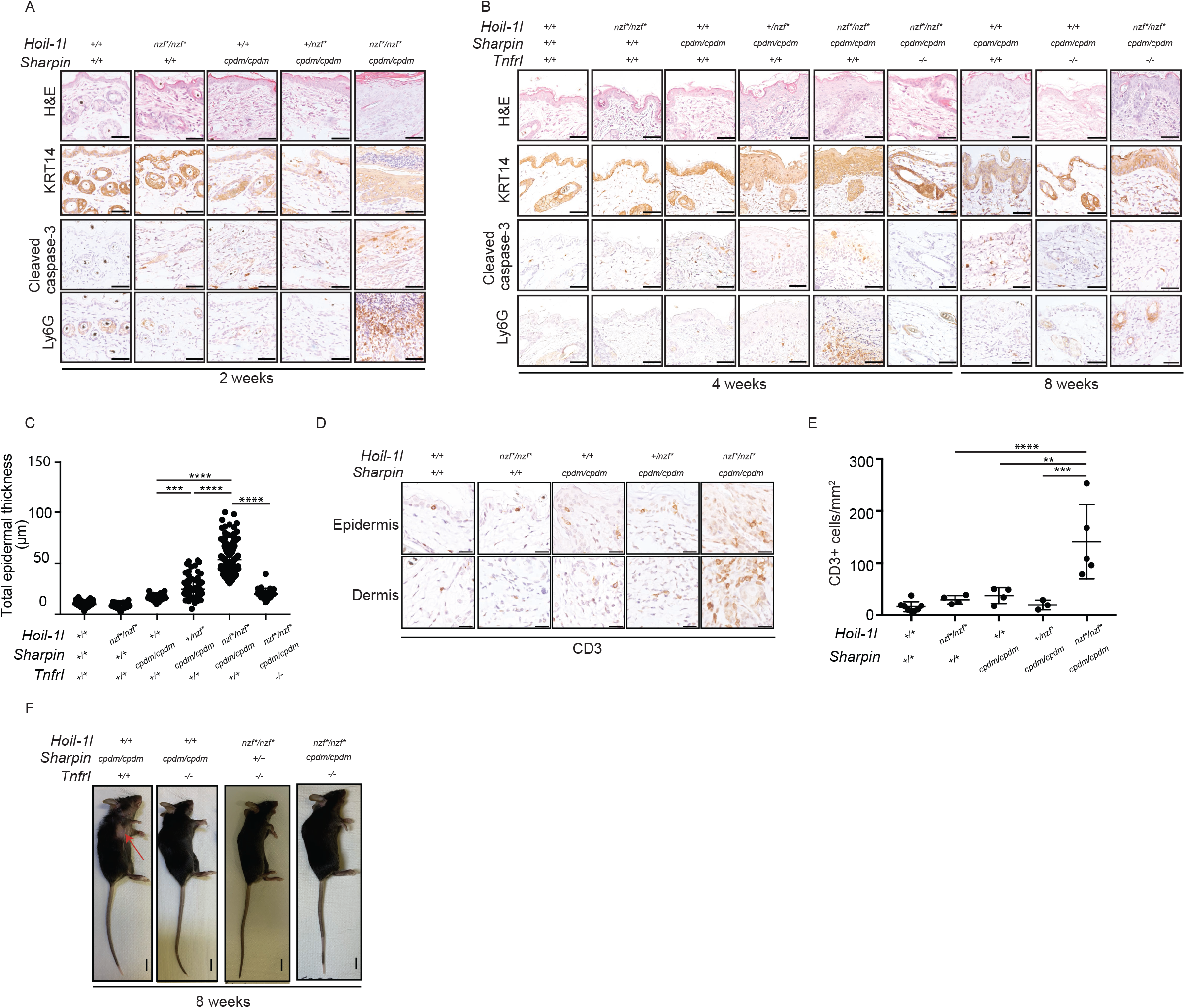
TNFR1 knockout resolves skin inflammation and keratinocyte apoptosis in *Hoil-1l*^*nzf*/nzf\**^; *Sharpin*^*cpdm/cpdm*^ mice. **A-B**. H&E staining, cleaved caspase-3, Keratin 14 (KRT14) and Ly6G immunostaining of ventral skin sections of 2, 4 or 8-week-old mice from the indicated genotypes. Scale bar: 50 μm. **C**.Quantification of the total epidermal thickness from KRT14-immunostained ventral skin sections of 4-week-old mice from the indicated genotypes. **D**.CD3 staining of epidermis and dermis of ventral skin sections from 4-week-old mice of the indicated genotypes. Scale bar: 25 μm. **E**.Quantification of CD3 signal in the ventral skin (epidermis and dermis) from 4-week-old mice of the indicated genotypes. **F**.Representative image of the gross appearance from 8-week-old mice of the indicated genotypes. The red arrow indicates skin lesions. **Data information**. Data are represented as mean ± SD, ANOVA, **p-value≤0.01, ***p-value≤0.001, ****p-value ≤0.0001. (C and F) *Hoil-1l*^*+/+*^; *Sharpin*^*+/+*^ (n=5 for C, n=8 for F), *Hoil-1l*^*nzf*/nzf\**^; *Sharpin*^*+/+*^ (n=4 for C and F), *Hoil-1l*^*+/+*^; *Sharpin*^*cpdm/cpdm*^ (n=4 for C and F), *Hoil-1l*^*+/nzf\**^; *Sharpin*^*cpdm/cpdm*^ (n=6 for C, n=3 for F), *Hoil-1l*^*nzf*/nzf\**^; *Sharpin*^*cpdm/cpdm*^ (n=5 for C and F). (C) *Hoil-1l*^*nzf*/nzf\**^; *Sharpin*^*cpdm/cpdm*^; *Tnfr1*^*-/-*^ (n=3).

Because the *Sharpin*^*cpdm/cpdm*^ skin phenotype is dependent on TNFR1 signaling (Gerlach *et al*., 2011; Kumari *et al*., 2014; Rickard *et al*., 2014), we hypothesized that the increased skin inflammation and apoptosis observed in *Hoil-1l*^*nzf*/nzf\**^; *Sharpin*^*cpdm/cpdm*^ mice are TNFR1-dependent. To address this, we generated TNFR1-deficient mice in the background of *Hoil-1l*^*nzf*/nzf\**^; *Sharpin*^*cpdm/cpdm*^ and analyzed their phenotype. Consistent with our hypothesis, TNFR1 knockout mitigated the skin inflammation and apoptosis observed in the 4-week-old *Hoil-1l*^*nzf*/nzf\**^; *Sharpin*^*cpdm/cpdm*^ mice (Figure 6B-D). However, at a later time point, 8 weeks, these triple mutant mice started to display mild signs of skin inflammation and apoptosis, suggesting the TNFR1-pathway is not the only pathway that regulates skin inflammation in these mice (Figure 6B and 6F). The TNFR1 knockout resolved additional phenotypes of *Hoil-1l*^*nzf*/nzf\**^; *Sharpin*^*cpdm/cpdm*^ mice, including body weight, spleen weight, and spleen pathology (Figure 5C-F), indicating an involvement of the TNFR signaling pathway.

Collectively, the linear ubiquitin chain binding domain of HOIL-1L cooperates with SHARPIN to restrict excessive skin inflammation and cell death, which is at least partially dependent on TNFR1. Furthermore, an imbalance in the population of immune cells could contribute to the inflammatory phenotype observed in *Hoil-1l*^*nzf*/nzf\**^; *Sharpin*^*cpdm/cpdm*^ mice.

## Discussion

Progress towards understanding the function of the LUBAC component HOIL-1L has been slow when compared to the other components, HOIP (a catalytic E3 ligase for linear ubiquitination) and SHARPIN (an adaptor protein). A crystal structure of a HOIL-1L fragment revealed that the NZF domain has a special C-terminal stretch that enables specific binding of linear ubiquitin chains (Sato *et al*., 2011). Among seven known linear ubiquitin specific binders (Fennell, Rahighi and Ikeda, 2018), HOIL-1L is the only component of the LUBAC complex that also generates linear ubiquitin chains. Yet the physiological function of this HOIL-1L activity *in vivo* had remained totally unknown. We previously showed that the linear ubiquitin chain-specific binding domain in NEMO (called UBAN) is required for NF-κB activation in cells (Rahighi *et al*., 2009).In this study, we uncovered that HOIL-1L NZF contributes the TNF-induced signaling pathway by promoting proper TNFR Complex I formation in cells. Furthermore, by generating HOIL-1L NZF mutant knockin mice, we for the first-time revealed functions of the HOIL-1L NZF domain in immune responses *in vivo*.

In general, it is known that the TNF-dependent NF-κB signaling pathway contributes to LPS-induced septic shock by promoting pro-inflammatory cytokine induction (Mandal *et al*., 2018; Vandewalle *et al*., 2019). When the NF-κB pathway is blocked, target cytokine induction *in vivo* is reduced, immune responses are suppressed, and mice are more resistant to TNF- and LPS-induced shock (Sheehan, Ruddle and Schreiber, 1989; Marino *et al*., 1997; Tortola *et al*., 2016; Mandal *et al*., 2018; Vandewalle *et al*., 2019). In the case of *Hoil-1l*^*nzf*/nzf\**^ mice, LPS-induced and TNF-induced shock were both milder than control mice, consistent with their reduced TNF-induced NF-κB target gene expression. In macrophages, we observed similar levels of NF-κB-dependent gene induction by the LSP-TLR4 pathway in wild-type and *Hoil-1l*^*nzf*/nzf\**^, thus we speculate that the resistance to LPS-induced septic shock is indirectly due to the diminished TNFR pathway (summarized in Figure 7). In addition to NF-κB signaling, linear ubiquitin chain binding by the HOIL-1L NZF might be required for the inflammasome-dependent pathway induced by septic shock. Inflammasome formation leads to the maturation and release of IL-1β, IL-18 and to pyroptosis causing tissue damage (Kayagaki *et al*., 2011; Broz and Dixit, 2016). Indeed, HOIL-1L was reported as a positive regulator of inflammasome assembly in cells (Rodgers *et al*., 2014). In support of this alternative, LPS-induced septic shock in *Hoil-1l*^*nzf*/nzf\**^ mice had lower serum levels of IL-1β serum than similarly treated wild type littermates. In the future, it will be important to decipher how HOIL-1L regulates inflammasome activation.

**Figure 7.**
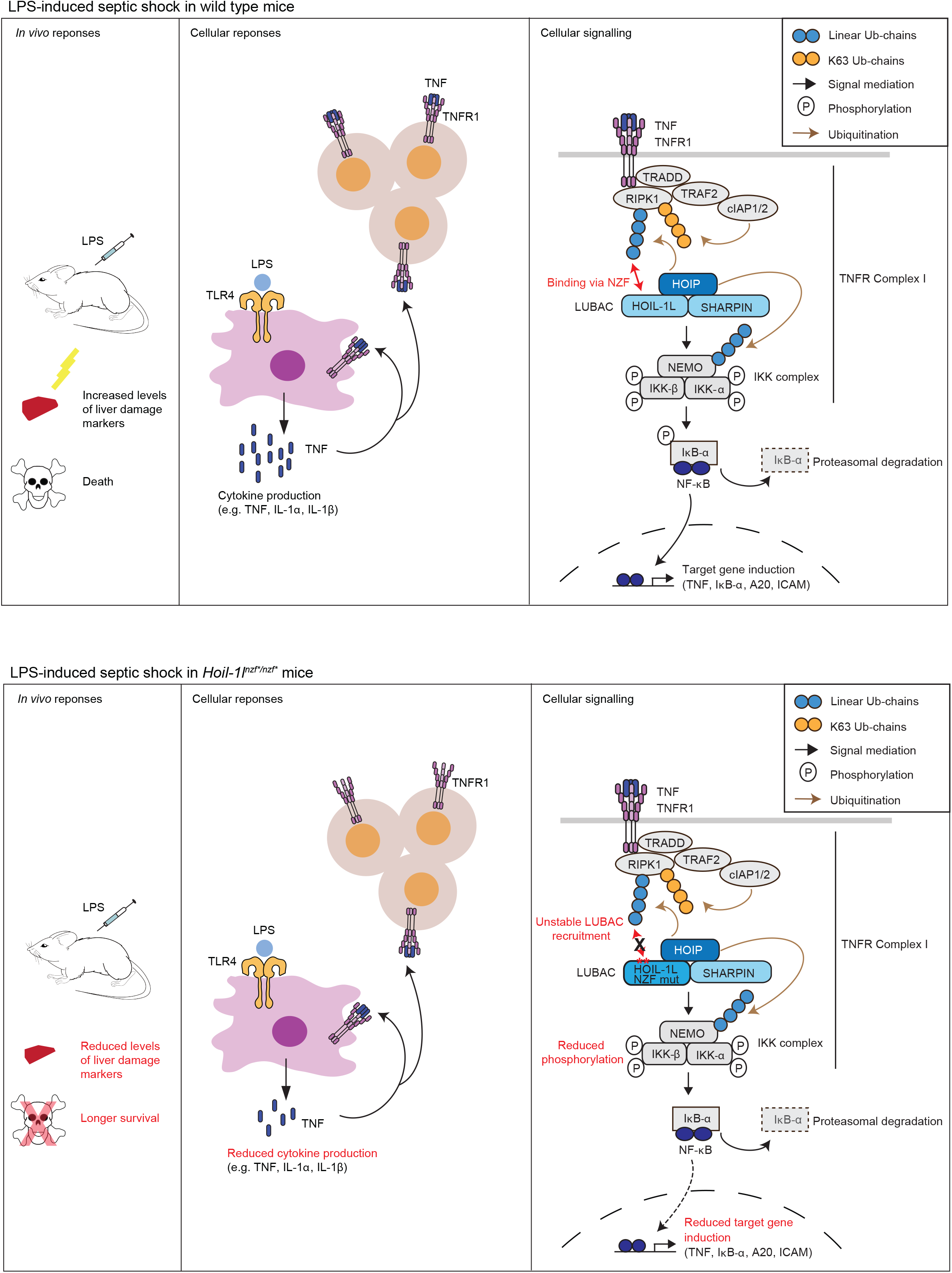
Proposed model depicting the role of the linear ubiquitin chain recognition by HOIL-1L-NZF in the regulation of the LPS-induced responses via the TNFR pathway. In wild type mice, LPS challenge leads to cytokine production, increased levels of liver damage markers, and death. At the cellular level, TNF-dependent NF-κB activation is mediated through LUBAC in which HOIL-1L NZF binds to linear ubiquitin chains to form TNFR complex I. A double point mutation in the HOIL-1L-NZF domain, abrogating interaction with linear ubiquitin chains, confers resistance to LPS-induced shock in mice, in which cytokine production is reduced. In cells from these mutant mice, recruitment of the LUBAC components in the TNFR complex I is more transient than wild type thus leading to reduced NF-κB activation.

The responses to LPS- or TNF-induced shock vary in viable mice with LUBAC disruptions, including *Sharpin*^*cpdm/cpdm*^ (Nastase *et al*., 2016), HOIL-1L partial deletion (originally described as knockout) (Tokunaga *et al*., 2009) and HOIL-1L ΔRING1 mice (Fuseya *et al*., 2020). *Sharpin*^*cpdm/cpdm*^ mice have severe LPS-induced septic shock which is dependent on the caspase-1 pathway (Nastase *et al*., 2016). TNF-induced shock in HOIL-1L partial deletion mice leads to liver apoptosis (Tokunaga *et al*., 2009). HOIL-1L ΔRING1 mice are more resistant to LPS and D-GalN-induced shock with reduced apoptosis in the liver (Fuseya *et al*., 2020). These data further indicate that LUBAC is involved in multiple pathways that control immune responses *in vivo*.

Systemic inflammation of *Sharpin*^*cpdm/cpdm*^ mice can be resolved by different genetic modifications; for example, TNF KO (Gerlach *et al*., 2011), TNFR1 or TRADD KO (Kumari *et al*., 2014), and caspase-8 KO or MLKL KO (Rickard *et al*., 2014) largely resolve the inflammatory phenotype, indicating that TNF-induced cell death drives inflammation in *Sharpin*^*cpdm/cpdm*^ mice. Furthermore, caspase-1 KO (Nastase *et al*., 2016), caspase 1/11 double KO (Douglas *et al*., 2015) also resolve the skin inflammation phenotype, suggesting that the caspase-1 pathway also plays an important role. More recently, HOIL-1L ΔRING1 mice were shown to resolve the *Sharpin*^*cpdm/cpdm*^ phenotype (Fuseya *et al*., 2020) whereas crosses with HOIL-1L ΔRBR mice (Shimizu *et al*., 2016) or HOIP mutant knockin mice of a specific ubiquitination site (Fennell *et al*., 2020) leads to embryonic lethality, suggesting a fine balance of ubiquitination processes regulated by LUBAC contributes to the *Sharpin*^*cpdm/cpdm*^ phenotype. Disruption of the HOIL-1L NZF domain exacerbated the skin inflammation phenotype of *Sharpin*^*cpdm/cpdm*^ mice, which was accompanied with keratinocyte apoptosis. Because TNFR1 KO largely resolved skin inflammation and apoptosis in these mice, we speculate that the HOIL-1L NZF cooperates with SHARPIN to prevent inflammation.

Additionally, we observed that the immune cell environment was altered by mutation of the HOIL-1L NZF domain in SHARPIN-deficient mice. *Sharpin*^*cpdm/cpdm*^ mice display splenomegaly and an increased percentage of neutrophils in the spleen, indicating that an imbalanced immune cell composition precedes the onset of dermatitis (Gurung, Sharma and Kanneganti, 2016). Interestingly, we observed a reduction of neutrophils in the spleens of *Hoil-1l*^*nzf*/nzf\**^; *Sharpin*^*cpdm/cpdm*^, despite more severe presentation of chronic proliferative dermatitis. On the other hand, *Hoil-1l*^*nzf*/nzf\**^; *Sharpin*^*cpdm/cpdm*^ displayed an increased frequency of plasma cells in the mesenteric lymph nodes, suggesting that these cells may contribute to the exacerbated skin inflammatory phenotype.

In conclusion, we demonstrate for the first time that the linear ubiquitin binding domain of HOIL-1L positively regulates the TNF-induced NF-κB signaling pathway both *in vitro* and *in vivo* (Figure 7). We propose that small molecules disrupting the interaction of HOIL-1L-NZF with linear ubiquitin chains may constitute a promising treatment for patients suffering from pathogen-associated septic shock.

## Material and Methods

### Antibodies and reagents

The antibodies used for immunoblotting and histological staining in this study are the following: anti-Myc (9E10) (Covance, MMS-150P), anti-Flag (M2) (Sigma, F3165), anti-vinculin (Sigma-Aldrich, V9131), anti-Tubulin (Abcam, ab15246), anti-ubiquitin (P4D1) (Santa Cruz Biotechnology, sc-8017), anti-HOIL-1L (Merck Millipore, MABC576, and Atlas Antibody, HPA 024185 used for detecting HOIL-1L in TNFR Complex I), anti-SHARPIN (Novus, NBP2-04116), anti-linear ubiquitin (LUB9) (Life Sensors, #AB130), anti-human HOIP (Sigma, SAB2102031), anti-mouse HOIP (homemade, described in (Fennell *et al*., 2020)), anti-IκB-α (Cell Signaling, #4812), anti-pIκB-α (Cell Signaling, #9246), anti-phosphorylated IκB-α (Cell Signaling, #9246), anti-IKK-α/β (Abcam, EPR16628), anti-IKK-α (CST2682, Bioké), anti-NEMO/IKKγ (FL-419) (Santa Cruz, sc-8330), anti-PARP (Cell Signaling, #9542), anti-cleaved caspase-3 (Cell Signaling, #9664), anti-FADD (for immunoprecipitation) (Santa Cruz, sc-271748), anti-FADD (for detection in cell lysates) (Abcam, ab124812), anti-RIPK1 (Cell signaling, #3493), IgG-HRP (Bio-Rad, 170-6516), goat anti-Rabbit IgG-HRP (Dako, P0448). Immunoprecipitations were performed using Protein G Agarose beads (Roche, 1124323301) or anti-FLAG (M2) affinity agarose gel (Sigma Aldrich, A2220). Other reagents used in this study are the following: human Flag-TNF (Enzo, ALX-522-008-C050), mouse TNF (Immunotools, 12343017), cycloheximide (CHX) (Sigma Aldrich, C4859), Z-Val-Ala-DL-Asp-fluoromethylketone (z-VAD-OMe-FMK) (Bachem, N-1560), Lipopolysaccharide (LPS) (from Escherichia coli O11:B4, L4391, Thermo Fisher).

### Plasmids

The following plasmids were used in previous studies: pBABE-puro-Flag-human SHARPIN (Ikeda *et al*., 2011), pEGFP-C1-human SHARPIN (Ikeda *et al*., 2011), pGEX-6P-1-human HOIP (Ikeda *et al*., 2011), pGEX-6P-1-human NEMO (Fennell *et al*., 2020), pGEX-6P-1-human HOIL-1L (Ikeda *et al*., 2011), pGEX-6P-1-human SHARPIN (Ikeda *et al*., 2011), pcDNA3-human-Ubiquitin (Ikeda *et al*., 2011), pGEX-4T1-diUbiquitin (Rahighi *et al*., 2009), pGEX-4T1-Empty (Rahighi *et al*., 2009), pcDNA3-human-HOIL-1L-HA (Tokunaga *et al*., 2009), pcDNA3-Myc-human HOIP (Tokunaga *et al*., 2009), pGex6P-1-human UbcH7 (Stieglitz *et al*., 2012) and pET49b-human HOIL-1L (C460A) (Stieglitz *et al*., 2012). The plasmids containing the point mutation in HOIP C885A was introduced into pcDNA3-Myc-human HOIP using site directed mutagenesis. Likewise, the point mutations in HOIL-1L T203A/R210A, and HOIL-1L C460A and the HOIL-1L deletion constructs ΔNZF and NZF-RBR were generated using pcDNA3-human-HOIL-1L-HA as template, following a standard protocol of site directed mutagenesis. The sequences of all plasmids were confirmed by Sanger sequencing. The sequence of the primers used for site-directed mutagenesis in this study are listed below.

HOIP C885A forward primer: 5’-GCCCGAGGAGGCGCCATGCACTTTCACTGTACC-3’, HOIP C885A reverse primer : 5’-GGTACAGTGAAAGTGCATGGCGCCTCCTCGGG-3’, HOIL-1L T203A forward primer : 5’-CAGTGCCCCGGGTGCGCCTTCATCAACAAGC-3’, HOIL-1L T203A reverse primer : 5’-GCTTGTTGATGAAGGCGCACCCGGGGCACTG-3’, HOIL-1L R210A forward primer : 5’-CAACAAGCCCACGGCGCCTGGCTGTGAG-3’, HOIL-1L R210A reverse primer: 5’-ACAGCCAGGCGCCGTGGGCTTGTTGATG -3’, HOIL-1L delta NZF forward primer: 5’ CGTCGGTCTACGTCACGGACTCGGGGGT-3’, HOIL-1L delta NZF reverse primer: 5’ TCAAGGGGAGGACGACGACGACGAAGGC-3’, HOIL-1L NZF+RBR forward primer: 5’CTTGAATTCATGGTGGGCTGGCAG 3’, HOIL-1L NZF+RBR reverse primer: 5’ CTGCCAGCCCACCATGAATTCAAG 3’

### Tissue culture and transfection

HEK293T (ATCC), primary and immortalized MEF and primary BMDM were cultured in Dulbecco’s modified Eagle’s medium (DMEM) (Sigma, D5648) supplemented with fetal calf serum (Thermo Fisher Scientific, 10270106), L-glutamine (Thermo Fisher Scientific, 25030-024), and penicillin–streptomycin (Sigma, P0781) and kept at 37°C and 5% CO2. Transfection of plasmids in MEF and HEK293T was carried out using GeneJuice Transfection Reagent (Merck Millipore, 70967) following the manufacturer’s protocol.

### NF-κB reporter assay

Method is described elsewhere (Ikeda *et al*., 2011; Fennell *et al*., 2020). Briefly, HEK293T were seeded in a 96-well plate and transfected with pNF-κB-Luc (Stratagene) and phRL-TK (Promega) as well as various HOIP, HOIL-1L and SHARPIN plasmids (as indicated) using GeneJuice. 48 hours post transfection, a Dual-glow luciferase assay (Promega E2940) was performed following the manufacturer’s protocol. Synergy H1 hybrid multimode microplate reader (BioTek) was used to measure the luminescence signal. The luciferase signal from each sample was normalized to the corresponding Renilla luciferase signal in each well. All samples were normalized to the control samples transfected with an empty vector (pcDNA3.1-Myc), pNF-κB-Luc and phRK-TK.

### Cell lysis and immunoprecipitation

Cells were lysed in lysis buffer containing 50 mM HEPES (pH7.4) (Sigma Aldrich, H4034), 150 mM NaCl, 1 mM EDTA, 1 mM EGTA, 1% Triton X-100, 10% Glycerol, 25 mM NAF and 10 μM ZnCl2 which was supplemented with 1 x cOmplete protease inhibitor cocktail (Roche, 11836170001), 1 mM PMSF (Roche, 10837091001) and 10 mM NEM (Sigma-Aldrich, E3876) (Fennell *et al*., 2020). Immediately after lysis, cells were centrifuged at 15,000 rpm for 15 minutes. Next the supernatant was denatured by adding SDS sample buffer and incubating at 96°C for 3 minutes. Cells lysates were incubated with 1μg of anti-Myc antibody for 2 hours at 4°C, followed by incubation with Protein G beads (Roche, 1124323301) for 2 hours at 4°C. Beads were washed 5-7 times in lysis buffer prior to elution with 30 μl of SDS sample buffer and heated for 3 minutes at 96°C.

### SDS-PAGE and immunoblotting

A method for immunoblotting is described before (Ikeda *et al*., 2011; Kumari *et al*., 2014). Briefly, samples were resolved by SDS-PAGE and subsequently transferred to a 0.45 μm nitrocellulose membrane (GE Healthcare, 10600019). The membrane was stained with Ponceau S to assess transfer efficiency. Next, membranes were washed with TBS-Triton X and TBS buffer and incubated in 5% BSA/TBS. Then membranes were incubated with the indicated primary antibodies diluted in 5% BSA/TBS overnight at 4°C. Membranes were incubated with the secondary antibody for 1-2 hours following the manufacturer’s recommendations. The signal was detected using Western Blot Luminol Reagent (Santa Cruz, sc-2048) on high performance chemiluminescence films (GE Healthcare, Amersham Hyperfilm ECL 28-9068-37)

### Protein purification

A method is described elsewhere (Ikeda *et al*., 2011; Asaoka *et al*., 2016). Briefly, expression plasmids were transformed into BL21 (DE3) *Escherichia coli*. Bacteria were grown at 37°C in LB media. The expression of GST-tagged fusion proteins was induced by adding IPTG 100 μM (Thermo Fischer Scientific, R0392) at OD_600_=0.8 in 4 to 6 liters of culture. To induce the expression of HOIP, HOIL-1L and SHARPIN, 100 μM ZnCl_2_ (Sigma Aldrich, 229997) were added. Bacteria cultures were grown overnight at 18°C. Bacteria were centrifuged, pelleted and resuspended in a buffer containing 100 mM HEPES (Sigma Aldrich, H4034), 600 mM NaCl, 1 mM TCEP-HCl (Thermo Fisher Scientific 20491), pH 7.4, supplemented with recombinant DNASE I (1000 U) (Roche, 0453628001), cOmplete protease EDTA-free inhibitor cocktail (Roche, 11836170001) and 1 mM PMSF (100mM in isopropanol, Roche 10837091001). Subsequently, bacteria were sonicated and 0.5% TritonX-100 was added to the lysate. The lysate was cleared by centrifugation and it was applied to a 5 ml GSTrap FF column (GE Healthcare, 17513101) to purify GST-proteins. PreScission protease (homemade) was used to remove the GST tag. Size exclusion chromatography on gel filtration columns using either the Superdex 200 (16/600) (GE Healthcare, GE28-9893-35) or Superdex 75 (16/600) (GE Healthcare, GE28-9893-33) in a buffer containing 50 mM HEPES (Sigma Aldrich, H4034), 150 mM NaCl, 1 mM TCEP-HCl (Thermo Fisher Scientific, 20491), pH 7.4 was used to resolve the protein eluates. The eluted fractions were subjected to SDS page and stained with InstantBlue™ and the fractions containing the desired proteins were pooled together. The protein concentration was assessed by comparison to BSA standards. The baculovirus used for insect expression of His_6_ mouse Ube1 in Hi5 cells was purified as previously described (Iwai *et al*., 1999; Fennell *et al*., 2020). For GST-empty and GST-Liner di-Ubiquitin, plasmids were expressed in BL21 cells for 16 hours at 25°C and lysed by sonication in the buffer described above. Subsequently, the lysates were incubated 16 hours at 4°C with Glutathione Sepharose 4B (GE Healthcare, GE17-0756-01). Next, beads were washed in 50 mM Tris, 100 mM EDTA, 150 mM NaCl, 0.5% Triton, pH 7.5 and resuspended in a buffer containing 50mM Tris, pH 7.5.

### *In vitro* ubiquitination assay

A method for *in vitro* ubiquitination assays is described elsewhere (Asaoka *et al*., 2016; Fennell *et al*., 2020). Ubiquitin (10 µg) (Sigma-Aldrich, U6253), mouse E1 (Ube1) (150 ng), human Ubch7 (300 ng), human HOIP (5 µg), human SHARPIN (1µg), human HOIL-1L (1 µg), human NEMO (5 µg) and ATP (2 mM) (Roche, 10519979000) were incubated at 37°C for the indicated time in a buffer containing 150 mM NaCl, 20 mM MgCl_2_ and 50 mM HEPES (Sigma Aldrich, H4034) (pH7.5). Reactions were terminated by adding SDS sample buffer and incubation at 96°C for 1 minute. Samples were subjected to SDS-PAGE followed by immunoblotting.

### GST Pull-Down assay

A method is described elsewhere (Rahighi *et al*., 2009). Briefly, GST-empty and GST-Linear di ubiquitin were expressed in *Escherichia. coli*, purified, immobilized on glutathione Sepharose 4B beads (Millipore Sigma, GE 17-075601) and incubated with the recombinant proteins or total cell lysates for 12 hours at 4°C. Next, GST-pulldown samples were washed five times with ice-cold lysis buffer (for total cell lysate samples) or with wash buffer (for recombinant protein samples) (25 mM Tris-HCl buffer (pH 7.2), 20 μM ZnCl_2_, 1 mM DTT, 100 mM NaCl, and 0.1% Triton X-100) (Sato *et al*., 2011).

### Mouse lines and husbandry

C57BL/KaLawRij *Sharpin*^*cpdm/cpdm*^ mice were described elsewhere (Seymour *et al*., 2007) (JAX stock #007599). C57BL/6J *Tnfr1-/-*(*Tnfrsf1a*^*tm1Mak*^/*TNFRp55-deficient* (B6.129*-Tnfrsf1a*^*tm1Mak*^/*J*)) were described elsewhere (Pfeffer *et al*., 1993). Mice were housed under specific pathogen free (SPF) conditions in individually ventilated cages with a HEPA filtered air (TECNIPLAST Green line GM 500) in a 14 hour-light/10 hour-dark cycle. The microbiological status of the mouse colony was monitored by sentinel mice (solid bedding sentinels and contact sentinels). *Hoil-1l*^*nzf*/nzf\**^ mice were backcrossed with C57BL/6J mice and subsequently used to generate all the genotypes used in the study and to maintain the line. *Hoil-1l*^*+/nzf\**^ C57BL/6J female and male mice, or *Hoil-1l*^*nzf*/nzf\**^ C57BL/6J female and male mice were crossed with *Sharpin*^*+/cpdm*^ male and female mice, which were already backcrossed with C57BL/6J mice. *Hoil-1l*^*+/nzf*;*^ *Sharpin*^*+/cpdm*^ male and female mice were crossed to obtain all genotypes used in this study. To maintain this mouse line *Hoil-1l*^*+/nzf*;*^ *Sharpin*^*+/cpdm*^ male and female mice obtained from different litters were bred. *Hoil-1l*^*+/nzf*;*^ *Sharpin*^*+/cpdm*^ female mice were bred with *Tnfr1*^*-/-*^ male mice. *Hoil-1l*^*+/nzf\**^; *Sharpin*^*+/cpdm*^; *Tnfr1*^+/-^ male and female mice obtained from these crosses were bred to obtain all genotypes used in this study. To maintain this mouse line, *Hoil-1l*^*+/nzf*;*^ *Sharpin*^*+/cpdm*^; *Tnfr1*^+/-^ male and female mice and *Hoil-1l*^*+/nzf*^ ; *Sharpin*^*+/cpdm*^; *Tnfr1*^-/-^ male and female mice obtained from different litters were crossed. All animal procedures described here were approved by the competent authorities.

### Generation of *Hoil-1l*^*nzf*/nzf\**^ mice

A method is described elsewhere (Wang *et al*., 2013). Briefly, the design of the gRNA was performed using the guide design tool from the Zhang lab (crispr.mit.edu). The gRNA template was generated using a forward and reverse oligonucleotide (IDT, HPLC purity) containing BbsI overhangs that were annealed and phosphorylated using T4 Polynucleotide kinase (PNK) (New England Biolabs, M0201S). Next, the px330 plasmid (Addgene plasmid number #42330, a gift from Feng Zhang (Cong *et al*., 2013) was digested with BbsI and dephosphorylated using calf intestinal alkaline phosphatase (CIP) (NEB, M0290S). The diluted phosphorylated oligonucleotide duplex was ligated into the dephosphorylated px330 plasmid using a T4 ligase (New England Biolabs, M0202M). Next, the ligated product was transformed into *Stbl3 E. coli* competent cells. The T7 promoter was added to the gRNA template by PCR amplification using primers that contain the T7 promoter sequence and gRNA sequence. The T7-gRNA PCR product was gel purified and used as the template for *in vitro* transcription (MEGA shortscript T7 kit, Invitrogen AM1345). The *in vitro* transcribed gRNA was then purified using the Invitrogen MEGAclear kit, AM1908) and eluted in non-DEPC RNAse free water (Ambion, AM9938). Cas9 mRNA was purchased from Sigma (CAS9MRNA-1EA). The single strand donor oligonucleotide contained the double point mutation in HOIL-1L-NZF (T201A/R208A), additionally it also contains a silent mutation of the PAM sequence and a silent mutation that generates a SmaI restriction site that allows genotyping of the mice. A scheme of the targeted strategy can be found in the results section. The primers, the sequence of the repair template and the single strand oligonucleotide donor template used for the generation of the *Hoil-1l*^*nzf*/nzf\**^ knockin mouse are listed below: gRNA sequence for BbsI overhang (forward primer): 5’ CACCGCTCACACCCAGGCCGTGT 3’ (gRNA A), 5’ CACCGTCTCACACCCAGGCCGTG 3’ (gRNA B). gRNA sequence for BbsI overhang (reverse primer): 5’ AAACACACGGCCTGGGTGTGAGC 3’ (gRNA A), 5’ AAACCACGGCCTGGGTGTGAGAC 3’ (gRNA B). Forward primer for *in vitro* transcription: 5’ TTAATACGACTCACTATAGGG 3’. Reverse primer for *in vitro* transcription: 5’ AAAAGCACCGACTCGGTGCC 3’. Single strand oligonucleotide donor template: 5’CCGGGCCCGGCTTTCATCAACAAACCTACTGCGCCTGGGTGTGAGATG 3’.C57BL/6J female donor mice (3-5-week-old) were treated with 5IU of pregnant mare’s serum gonadotropin (PMSG). 46 hours later the donor mice were injected with 5IU of human chorionic gonadotropin (hCG) (Intervet, GesmbH) to induce super ovulation. Next the females were mated with stud males and checked for plugs. The zygotes were isolated from pregnant females in M2 media and culture in KSOM media. Next a mix consisting of 100 ng/ μl of Cas9 mRNA, 50 ng/μl of gRNA and 200 ng/μl of single stand oligonucleotide diluted in non-DEPC RNase free water (Ambion, AM938) was centrifuged for 1 hour and 30 minutes at 15,000 rpm at 4°C and injected into the cytosol of zygotes (Volume of injection mixture: 50 μl). Following the injection, the zygotes were transferred to pseudo-pregnant females. 3 different founder lines obtained from two different gRNAs were established. The correct sequence for each founder mouse, containing the double point mutations and the silent mutations for genotyping purpose was confirmed by Sanger Sequencing. PCR fragments amplified from genomic DNA extracted from the different founder mice using the Wizard SV genomic DNA extraction kit (Promega, A2360), were cloned into a vector using the TOPO TA cloning kit (Thermo Fisher Scientific).

### Genotyping of *Hoil-1l*^*nzf*/nzf\**^ mice

Method is described in a previous study (Fennell *et al*., 2020). Briefly, genomic DNA from Proteinase K-digested mouse toes was extracted using the Wizard SV genomic DNA extraction kit (Promega, A2360). PCR reactions to amplify a fragment containing the target sites were carried out using the forward and reverse primers (Forward primer: 5’ TGGGTTGTCAGCATGTGGTT 3’. Reverse primer: 5’ GTGGTTCCCTTTCTGGCTCA 3’). PCR product was digested with SmaI (Thermo Fisher Scientific, ER0662) for 16 hours. Digested products were analyzed by electrophoresis using 1.5% agarose gels.

### Isolation and immortalization of Mouse Embryonic Fibroblasts (MEFs)

Primary MEFs were isolated from E13.5 embryos according to a standard protocol as described previously (Fennell *et al*., 2020). Briefly, female mice were crossed with a male mouse and checked daily for plugs. When a plug was detected, this was considered developmental stage E0.5. Embryos were isolated at E13.5. The head of the embryos was removed, and the rest of the body was minced with a scalpel and trypsinized for 5 minutes at 37°C. Cells were collected in a falcon tube, centrifuged and plated in a culture dish. To immortalize primary MEFs, pSG5-SV40 largeT antigen plasmid was transfected using GeneJuice Transfection Reagent following the manufacturer’s protocol and cells were kept until they became stably proliferative.

### Isolation and stimulation of Bone Marrow Derived Macrophages (BMDMs)

A method is described elsewhere (Baccarini, Bistoni and Lohmann-Matthes, 1985). Briefly, the bone marrow cells of the tibia and femur of 8 to 12-week-old mice (female and male) were flushed out and differentiated by culturing in DMEM-10% FCS supplemented with mouse colony stimulating factor (M-CSF, 25ng/ml, Peprotech, 315-02) for 5-6 days. For signaling assays, 1×10^6^ BMDM were seeded in 6 well plates, and stimulated with mouse TNF (20ng/ml) after 16 hours of serum starvation or stimulated with LPS (10ng/ml) for the indicated time.

### Isolation of TNFR Complex I

A method is previously described (Haas *et al*., 2009; Draber *et al*., 2015; Fennell *et al*., 2020). Briefly, 5-20×10^6^ MEFs were seeded in 15 cm dishes. Following 16-hour serum starvation in 0.2% FCS-DMEM, MEFs were treated with 1µg/ml of recombinant Flag-human TNF. Subsequently, cells were washed twice with chilled PBS and lysed in IP-lysis buffer containing (30 mM Tris-HCI (pH 7.4), 120 mM NaCl, 2 mM EDTA, 2 mM KCI, 10% glycerol, 1%Trition X-100, 50 mM NaF, 1 x cOmplete protease inhibitor cocktail (Roche, 11836170001), 1 mM PMSF (Roche, 10837091001), 10 mM NEM (Sigma-Aldrich, E3876), and 5 mM Na_3_VO_4_ (Sigma-Aldrich, S6508). Samples were incubated on ice for 30 minutes. Following centrifugation at 15,000rpms for 30 minutes, Flag human TNF (1 µg) was added to the 0-hour control samples. Samples were precleared with Protein G agarose beads for 1 hour at 4°C. anti-FLAG M2 beads (Sigma Aldrich, A2220) were incubated with the pre-cleared samples at 4°C. Samples were washed 6 times with IP-lysis buffer. Samples were eluted by incubation with 2X SDS sample buffer at 96°C for five minutes.

### qRT-PCR

This method is described elsewhere (Asaoka *et al*., 2016; Fennell *et al*., 2020). 1×10^6^ primary BMDMs or 1×10^6^ MEFs were serum starved in 0.2% FBS-DMEM for 15-16 hours and then treated with mouse TNF (20 ng/ml) for the indicated times. Samples were washed with chilled PBS and RNA was extracted using TRIzol (Life Technologies, 15596018) and treated with TURBO DNA-free kit (Invitrogen, AM1907). cDNA was generated using oligo (dT) 18 primer (New England Biolabs, 513165) and SuperScript II Reverse Transcriptase (Invitrogen, 18064-014) following the manufacturer’s protocol. Real time quantitative PCR was performed in a CFX 96 BioRad CFX 96 Real-Time PCR detection instrument with GoTaq qRT-PCR master mix (Promega, A6002). The sequence of the primers used in this study are: β-actin forward primer 5’-CGGTTCCGATGCCCTGAGGCTCTT-3’, β-actin reverse primer 5’ CGT CACACTTCATGATGGAATTGA-3. A20: forward primer 5’ AAAGGACTACAGCAGAGCCCAG-3’, A20 reverse primer 5’-AGAGACATTTCCAGTCCGGTGG-3’. ICAM forward primer 5’-AAGGAGATCACATTCACGGTG-3’, ICAM reverse primer 5’-TTTGG GATGGTAGCTGGAAG-3’, IκB-α forward primer 5’-GCTGAGGCACTTCTGAAAGCTG-3’, IκB-α reverse primer 5’-TGGACTGGCAGACCTACCATTG-3’. VCAM forward primer 5’-CTGGGAAGCTGGAACGAAGT-3’, VCAM reverse primer 5’-GCCAACACTTGACC GTGAC-3’. mouse TNF forward primer 5’ CATCTTCTCAAAATTCGAGTGACAA, mouse TNF reverse primer 5 ‘ TGGGAGTAGACAAGGTACAACCC. IL-6 forward primer 5’-AGCCAG AGTCCTTCAGAGAGA-3’, IL-6 reverse primer 5’-TGGTCTTGGTCCTTAGCCAC-3’. IL1-β forward primer 5’-ATGAAAGACGGCACACCCAC-3’, IL1β reverse primer 5’-CTGCTTGTGAGGTGCTGATG-3’CCL5 forward primer 5-TGC TGCT TTGCCTACCTC TC-3’, CCL5 reverse primer 5’-CCA C TTCTTCTCTGGGTTGG-3’.

### Caspase-8 activity assay

A method is described elsewhere (Kumari *et al*., 2014; Fennell *et al*., 2020). Briefly, MEFs were seeded at a density of 5×10^4^ cells/well in a 96-well white plate (Thermo Fisher Scientific, 136101) and treated with mouse TNF (100 ng/ml) and cycloheximide (1 µg/ml) for the indicated timepoints. Caspase Glow 8 assay system (Promega, G8202) was used to measure caspase-8 activity following the manufacturer’s instructions. Synergy H1 microplate reader (Fisher Scientific) was used to measure the signal.

### Flow cytometry

The spleen and mesenteric lymph nodes were harvested from 4-week-old mice. The peritoneal exudate was harvested as descried below. Spleen homogenates were lysed with Ammonium Chloride Potassium (ACK) lysis buffer to lyse red blood cells. Subsequently, cells were stained with the viability dye eFluor780 (eBioscience, 65-0865-14) at 4°C for 15 minutes. Cells were washed and blocked with CD16/CD32 Fc block (BD Bioscience, 553141) at 4°C for 10 minutes. The following antibodies were used for staining: CD23 (eBioscience, 25-0232-82), CD93 (Invitrogen, 62-5892-82), CD19 (BD Biosciences, 563333), CD45 (BioLegend, 103149), CD138 (BD Biosciences, 562935), TCRβ (BD Biosciences,109229), CD11c (BioLegend, 117335), MHC II (BioLegend, 107608), CD11b (BioLegend, 101257), TCRβ (BioLegend, 109205), CD23 (Biolegend, 101620), DX5 (BD Biosciences, 558295), F4/80 (BioLegend, 123131), B220 (BD Biosciences, 552772), CD8 (BD Biosciences, 553035), NK1.1 (BD Biosciences, 553164), TCRβ (BD Biosciences, 560729), CD23 (BioLegend, 101614), CD21 (BioLegend, 123411), Ly6G (BioLegend, 127618), Ly6C (BD Biosciences, 560595), TCRβ (BD Biosciences, 553170), B220 (BD Biosciences, 562922), CD28 (BioLegend, 122010), CD4 (BioLegend, 100443), CD45 (BD Biosciences, 550994). The samples were acquired using an LSR Fortessa flow cytometer (BD). The data was analyzed using the FlowJo software.

### Peritoneal lavage

Mice were deeply anesthetized with 150 μg/g ketamine (Ketamidor, Richter pharma) and 15 μg/g xylazine (Sedaxylan, Dechra). Peritoneal cavity was cut open and the peritoneum was flushed with 7-8 ml of harvest solution (PBS, 0,02% EDTA). The peritoneal lavage was retrieved and kept on ice until further processing.

### TNF-induced shock

8 to10-week-old mice were injected intravenously (i.v.) via the tail vein with 450 µg/kg of endotoxin-free recombinant mouse TNF (ImmunoTools) in 200 µl of PBS (pH 6.8) (Tortola *et al*., 2016). Mouse body temperature was recorded with an infrared thermometer (BIO-IRB153, Bioseb) by placing it 2-3 cm from the rectum (Mei *et al*., 2018). The weight, of the mice was monitored for 12 hours.

### LPS-induced septic shock

8 to 10-week old mice were injected intraperitoneally (i.p.) with 30 mg/kg of *E. coli* 0111: B4 LPS (L4391, Sigma) dissolved in 0.9% NaCl (Lamkanfi *et al*., 2009). Mice body temperature was recorded with an infrared thermometer (BIO-IRB153,Bioseb) by placing it 2-3 cm from the rectum (Mei *et al*., 2018). The weight and survival of the mice were monitored after the LPS-injection.

### AST/ALT measurements

12 hours post i.p. LPS injection (30 mg/kg), mice were deeply anesthetized with 150 μg/g ketamine (Ketamidor, Richter pharma) and 15 μg/g xylazine (Sedaxylan, Dechra). Blood was withdrawn from the vena cava with a 25-gouge heparinized needle connected to a syringe containing 50 μl of 100 U/mL heparin (Sigma-Aldrich) to prevent coagulation and the total final volume was noted. Subsequently, blood was centrifuged at 12000xg for 5 minutes and the serum was collected. The concentration of liver damage marker AST and ALT were determined by the veterinary *in vitro* diagnostics laboratory InVitro GmbH and normalized to the volume of withdrawn blood.

### Cytokine analysis

To measure cytokine levels in the serum of mice, ProcartaPlex Immunoassays (Thermo Fisher Scientific) were used. For mice subjected to LPS-induced septic shock, blood was withdrawn as described in the AST/ALT measurements. For mice subjected to TNF-induced shock, blood was withdrawn from the submandibular vein using a lancet and collected in a BD Microtainer blood collection tube to avoid coagulation. Subsequently, blood was centrifuged at 6000xg for 3 minutes and plasma was collected. Samples were measured in duplicates following the manufacturer’s instructions. The fluorescence intensity of each sample was read in a Luminex analyzer.

### Histopathological analysis

A method is described elsewhere (Kumari *et al*., 2014). Briefly, mouse tissues were fixed in 10% neutral buffered formalin (Sigma, HT501128) for 12-16 hours, embedded in paraffin and sectioned for hematoxylin and eosin (H&E) staining using an automated stainer (Microm HMS 740). An automated stainer (Bond III, Leica) was used for immunohistochemistry. Primary antibodies for immunohistochemistry are keratin 14 (KRT14) (Sigma Aldrich, SAB4501657, 1:200) cleaved caspase-3 (Cell Signaling, 9661, 1:100), Ly6G (Abcam, ab2557, 1:500), CD3 (Abcam ab49943, 1:200) and B220/CD45R (BD Biosciences 550286, 1:100). The secondary antibodies used are goat anti-rabbit IgG (Dako, E0432 1:500), anti-rat IgG (Abcam, ab6733, 1:500). Signal was detected using the Leica Bond Intense R Detection system. A board-certified veterinary comparative pathologist evaluated the slides using a Zeiss Axioscope 2 MOT microscope and images were acquired with a SPOT Insight color camera (SPOT Imaging, Diagnostic Instrument, Inc.).

For skin thickness measurements, whole slide images of KRT14-stained sections were evaluated using the Panoramic Viewer software. From each digital scene spanning 4 mm, 10 non-contiguous, representative foci were selected, and the thickness of the entire epidermis was measured in each focus. For quantification of CD3 positive cells in the skin, QuPath v 0.2.3, an open source image analysis tool was used (Bankhead *et al*., 2017). Annotations of skin sections were generated with a pixel classifier after detecting the tissue with simple thresholder for the hematoxylin channel. The annotations were manually verified and refined by a pathologist. Positive cell detection was performed by setting the optical density sum for the Cell DAB OD mean score compartment. The threshold was applied to all images within the workspace with a command history script.

### Statistical analysis

Statistical analysis was performed using Prism 8 software (GraphPad). The mean values with standard deviation are shown. Parametric T-test was used for normally distributed datasets. ANOVA was used to analyze repeated measurements over time. The significance and confidence level were set at 0.05, and P values are indicated in each figure legends as well as the number of replicates used to calculate statistics.

## Acknowledgements

We thank all the members of the Ikeda lab, especially Alan Rodríguez Carvajal, and the Penninger lab for their support and constructive discussions on the project. We thank molecular biology services, bio-optics, bioinformatics, transgenic services (the IMP-IMBA core facilities), the Histopathology facility and the Protein Technologies facility (part of the Vienna Biocenter Core Facilities) for their technical support. We thank Margit Jaschke and Melanie Gierer (Thermo Fisher Scientific) for their support with the ELISA measurement and René Rauschmeier (IMP) for i.v. injections. J.M.P. is supported by the Austrian Federal Ministry of Education, Science and Research, the Austrian Academy of Sciences and the City of Vienna and grants from the Austrian Science Fund (FWF) Wittgenstein award (Z 271-B19), the T. von Zastrow foundation, a Canada 150 Research Chairs Program (F18-01336), and CIHR grant. Research in the Ikeda Lab is supported by JSPS KAKENHI Grant Number 18K19959, ERC Consolidator Grant (LUbi, 614711) and the Austrian Academy of Sciences. We also thank Angela Andersen from the Life Science Editors for editing the manuscript.

## Author contributions

C.G.D and F.I designed the experiments, prepared the figures and wrote the manuscript. C.G.D, G.J, K.S, L.D, A.B, K.E, J.A, and L.M.F performed experiments and analyzed data. A.K performed histopathology analysis. A.B, K.E, P.K and A.H contributed to *in vivo* experiments and provided essential technical advice. J.M.P, and F.I supervised author students and secured funding. F.I. conducted the overall project.

## Conflict of interests

The authors declare no conflict of interests.

## References

Asaoka, T. et al.. (2016) ‘Linear ubiquitination by LUBEL has a role in Drosophila heat stress response’, EMBO reports, 17(11), pp. 1624–1640. doi: 10.15252/embr.201642378.

Asaoka, T. and Ikeda, F. (2015) ‘New Insights into the Role of Ubiquitin Networks in the Regulation of Antiapoptosis Pathways’, International Review of Cell and Molecular Biology, 318, pp. 121–158. doi: 10.1016/bs.ircmb.2015.05.003.

Baccarini, M., Bistoni, F. and Lohmann-Matthes, M. L. (1985) ‘In vitro natural cell-mediated cytotoxicity against Candida albicans: macrophage precursors as effector cells’, Journal of Immunology (Baltimore, Md.: 1950), 134(4), pp. 2658–2665.

Bankhead, P. et al.. (2017) ‘QuPath: Open source software for digital pathology image analysis’, Scientific Reports, 7(1), p. 16878. doi: 10.1038/s41598-017-17204-5.

Boisson, B. et al.. (2012) ‘Immunodeficiency, autoinflammation and amylopectinosis in humans with inherited HOIL-1 and LUBAC deficiency’, Nature Immunology. doi: 10.1038/ni.2457.

Boisson, B. et al.. (2015) ‘Human HOIP and LUBAC deficiency underlies autoinflammation, immunodeficiency, amylopectinosis, and lymphangiectasia’, Journal of Experimental Medicine. doi: 10.1084/jem.20141130.

Broz, P. and Dixit, V. M. (2016) Inflammasomes: Mechanism of assembly, regulation and signalling, Nature Reviews Immunology. Nature Publishing Group (7). doi: 10.1038/nri.2016.58.

Carvajal, A. R. et al.. (2020) The linear ubiquitin chain assembly complex LUBAC generates heterotypic ubiquitin chains. preprint. Biochemistry. doi: 10.1101/2020.05.27.117952.

Cong, L. et al.. (2013) ‘Multiplex genome engineering using CRISPR/Cas systems’, Science, 339(6121), pp. 819–823. doi: 10.1126/science.1231143.

Douglas, T. et al.. (2015) ‘The Inflammatory Caspases-1 and -11 Mediate the Pathogenesis of Dermatitis in Sharpin-Deficient Mice’, The Journal of Immunology, 195(5), pp. 2365–2373. doi: 10.4049/jimmunol.1500542.

Draber, P. et al.. (2015) ‘LUBAC-Recruited CYLD and A20 Regulate Gene Activation and Cell Death by Exerting Opposing Effects on Linear Ubiquitin in Signaling Complexes’, Cell Reports, 13(10), pp. 2258–2272. doi: 10.1016/j.celrep.2015.11.009.

Elton, L. et al.. (2016) ‘MALT1 cleaves the E3 ubiquitin ligase HOIL-1 in activated T cells, generating a dominant negative inhibitor of LUBAC-induced NF-κB signaling’, FEBS Journal, 283(3), pp. 403–412. doi: 10.1111/febs.13597.

Fennell, L. M. et al.. (2020) ‘Site-specific ubiquitination of the E3 ligase HOIP regulates apoptosis and immune signaling’, The EMBO Journal, 39(24). doi: 10.15252/embj.2019103303.

Fennell, L. M., Rahighi, S. and Ikeda, F. (2018) ‘Linear ubiquitin chain-binding domains’, The FEBS Journal, 285(15), pp. 2746–2761. doi: 10.1111/febs.14478.

Fuseya, Y. et al.. (2020) ‘The HOIL-1L ligase modulates immune signalling and cell death via monoubiquitination of LUBAC’, Nature Cell Biology, 22(6), pp. 663–673. doi: 10.1038/s41556-020-0517-9.

Gerlach, B. et al.. (2011) ‘Linear ubiquitination prevents inflammation and regulates immune signalling’, Nature, 471(7340), pp. 591–596. doi: 10.1038/nature09816.

Gómez-Díaz, C. and Ikeda, F. (2019) ‘Roles of ubiquitin in autophagy and cell death’, Seminars in Cell & Developmental Biology, 93, pp. 125–135. doi: 10.1016/j.semcdb.2018.09.004.

Gurung, P., Sharma, B. R. and Kanneganti, T. D. (2016) ‘Distinct role of IL-1β in instigating disease in Sharpin cpdm mice’, Scientific Reports, 6. doi: 10.1038/srep36634.

Haas, T. L. et al.. (2009) ‘Recruitment of the Linear Ubiquitin Chain Assembly Complex Stabilizes the TNF-R1 Signaling Complex and Is Required for TNF-Mediated Gene Induction’, Molecular Cell, 36(5), pp. 831–844. doi: 10.1016/j.molcel.2009.10.013.

Ikeda, F. et al.. (2011) ‘SHARPIN forms a linear ubiquitin ligase complex regulating NF-κB activity and apoptosis’, Nature, 471(7340), pp. 637–641. doi: 10.1038/nature09814.

Iwai, K. et al.. (1999) ‘Identification of the von Hippel-Lindau tumor-suppressor protein as part of an active E3 ubiquitin ligase complex’, Proceedings of the National Academy of Sciences, 96(22), pp. 12436–12441. doi: 10.1073/pnas.96.22.12436.

Justus, S. J. and Ting, A. T. (2015) ‘Cloaked in ubiquitin, A killer hides in plain sight: The molecular regulation of RIPK1’, Immunological Reviews, 266(1), pp. 145–160. doi: 10.1111/imr.12304.

Karki, R. et al.. (2021) ‘Synergism of TNF-α and IFN-γ Triggers Inflammatory Cell Death, Tissue Damage, and Mortality in SARS-CoV-2 Infection and Cytokine Shock Syndromes’, Cell. doi: 10.1016/j.cell.2020.11.025.

Kayagaki, N. et al.. (2011) ‘Non-canonical inflammasome activation targets caspase-11’, Nature, 479(7371), pp. 117–121. doi: 10.1038/nature10558.

Kelsall, I. R. et al.. (2019) ‘The E3 ligase HOIL-1 catalyses ester bond formation between ubiquitin and components of the Myddosome in mammalian cells’, Proceedings of the National Academy of Sciences of the United States of America, 116(27), pp. 13293–13298. doi: 10.1073/pnas.1905873116.

Kumari, S. et al.. (2014) ‘Sharpin prevents skin inflammation by inhibiting TNFR1-induced keratinocyte apoptosis’, eLife. doi: 10.7554/eLife.03422.

Lamkanfi, M. et al.. (2009) ‘Caspase-7 deficiency protects from endotoxin-induced lymphocyte apoptosis and improves survival’, Blood, 113(12), pp. 2742–2745. doi: 10.1182/blood-2008-09-178038.

Mandal, P. et al.. (2018) ‘Caspase-8 Collaborates with Caspase-11 to Drive Tissue Damage and Execution of Endotoxic Shock’, Immunity, 49(1), pp. 42-55.e6. doi: 10.1016/j.immuni.2018.06.011.

Marino, M. W. et al.. (1997) ‘Characterization of tumor necrosis factor-deficient mice’, Proceedings of the National Academy of Sciences, 94(15), pp. 8093–8098. doi: 10.1073/pnas.94.15.8093.

Mei, J. et al.. (2018) ‘Body temperature measurement in mice during acute illness: implantable temperature transponder versus surface infrared thermometry’, Scientific Reports, 8(1), p. 3526. doi: 10.1038/s41598-018-22020-6.

Nastase, M.-V. et al.. (2016) ‘An Essential Role for SHARPIN in the Regulation of Caspase 1 Activity in Sepsis’, The American Journal of Pathology, 186(5), pp. 1206–1220. doi: 10.1016/j.ajpath.2015.12.026.

Pahl, H. L. (1999) ‘Activators and target genes of Rel/NF-κB transcription factors’, Oncogene, 18(49), pp. 6853–6866. doi: 10.1038/sj.onc.1203239.

Peltzer, N. et al.. (2014) ‘HOIP deficiency causes embryonic lethality by aberrant TNFR1-mediated endothelial cell death’, Cell Reports. doi: 10.1016/j.celrep.2014.08.066.

Peltzer, N. et al.. (2018) ‘LUBAC is essential for embryogenesis by preventing cell death and enabling haematopoiesis’, Nature, 557(7703), pp. 112–117. doi: 10.1038/s41586-018-0064-8.

Peltzer, N. and Walczak, H. (2019) Cell Death and Inflammation – A Vital but Dangerous Liaison, Trends in Immunology. doi: 10.1016/j.it.2019.03.006.

Pfeffer, K. et al.. (1993) ‘Mice deficient for the 55 kd tumor necrosis factor receptor are resistant to endotoxic shock, yet succumb to L. monocytogenes infection’, Cell, 73(3), pp. 457–467. doi: 10.1016/0092-8674(93)90134-C.

Rahighi, S. et al.. (2009) ‘Specific Recognition of Linear Ubiquitin Chains by NEMO Is Important for NF-κB Activation’, Cell, 136(6), pp. 1098–1109. doi: 10.1016/j.cell.2009.03.007.

Rickard, J. A. et al.. (2014) ‘TNFR1-dependent cell death drives infammation in Sharpin-defcient mice’, eLife, 2014(3). doi: 10.7554/eLife.03464.

Rodgers, M. A. et al.. (2014) ‘The linear ubiquitin assembly complex (LUBAC) is essential for NLRP3 inflammasome activation’, Journal of Experimental Medicine, 211(7), pp. 1333–1347. doi: 10.1084/jem.20132486.

Sasaki, K. et al.. (2019) ‘Modulation of autoimmune pathogenesis by T cell-triggered inflammatory cell death’, Nature Communications, 10(1). doi: 10.1038/s41467-019-11858-7.

Sasaki, K. and Iwai, K. (2015) ‘Roles of linear ubiquitinylation, a crucial regulator of NF-κB and cell death, in the immune system’, Immunological Reviews, 266(1), pp. 175–189. doi: 10.1111/imr.12308.

Sato, Y. et al.. (2011) ‘Specific recognition of linear ubiquitin chains by the Npl4 zinc finger (NZF) domain of the HOIL-1L subunit of the linear ubiquitin chain assembly complex’, Proceedings of the National Academy of Sciences of the United States of America, 108(51), pp. 20520–20525. doi: 10.1073/pnas.1109088108.

Seymour, R. E. et al.. (2007) ‘Spontaneous mutations in the mouse Sharpin gene result in multiorgan inflammation, immune system dysregulation and dermatitis’, Genes & Immunity, 8(5), pp. 416–421. doi: 10.1038/sj.gene.6364403.

Sharma, B. R. et al.. (2019) ‘Innate immune adaptor MyD88 deficiency prevents skin inflammation in SHARPIN-deficient mice’, Cell Death and Differentiation, 26(4), pp. 741–750. doi: 10.1038/s41418-018-0159-7.

Sheehan, K., Ruddle, N. and Schreiber, R. (1989) ‘Generation and characterization of hamster monoclonal antibodies that neutralize murine tumor necrosis factors.’, 142 no. 11 3884–3893.

Shimizu, S. et al.. (2016) ‘Differential Involvement of the Npl4 Zinc Finger Domains of SHARPIN and HOIL-1L in Linear Ubiquitin Chain Assembly Complex-Mediated Cell Death Protection’, Molecular and Cellular Biology, 36(10), pp. 1569–1583. doi: 10.1128/MCB.01049-15.

Stieglitz, B. et al.. (2012) ‘LUBAC synthesizes linear ubiquitin chains via a thioester intermediate’, EMBO Reports, 13(9), pp. 840–846. doi: 10.1038/embor.2012.105.

Tokunaga, F. et al.. (2009) ‘Involvement of linear polyubiquitylation of NEMO in NF-κB activation’, Nature Cell Biology, 11(2), pp. 123–132. doi: 10.1038/ncb1821.

Tokunaga, F. et al.. (2011) ‘SHARPIN is a component of the NF-κB-activating linear ubiquitin chain assembly complex’, Nature, 471(7340), pp. 633–636. doi: 10.1038/nature09815.

Tortola, L. et al.. (2016) ‘The Tumor Suppressor Hace1 Is a Critical Regulator of TNFR1-Mediated Cell Fate’, Cell Reports, 15(7), pp. 1481–1492. doi: 10.1016/j.celrep.2016.04.032.

Vandewalle, J. et al.. (2019) ‘A Study of Cecal Ligation and Puncture-Induced Sepsis in Tissue-Specific Tumor Necrosis Factor Receptor 1-Deficient Mice’, Frontiers in Immunology, 10, p. 2574. doi: 10.3389/fimmu.2019.02574.

Wang, H. et al.. (2013) ‘One-step generation of mice carrying mutations in multiple genes by CRISPR/cas-mediated genome engineering’, Cell, 153(4), pp. 910–918. doi: 10.1016/j.cell.2013.04.025.

Webster, J. D. and Vucic, D. (2020) The Balance of TNF Mediated Pathways Regulates Inflammatory Cell Death Signaling in Healthy and Diseased Tissues, Frontiers in Cell and Developmental Biology. Frontiers Media S.A. doi: 10.3389/fcell.2020.00365.

Witt, A. and Vucic, D. (2017) Diverse ubiquitin linkages regulate RIP kinases-mediated inflammatory and cell death signaling, Cell Death and Differentiation. Nature Publishing Group (7). doi: 10.1038/cdd.2017.33.

